# Discovery of orally bioavailable SARS-CoV-2 papain-like protease inhibitor as a potential treatment for COVID-19

**DOI:** 10.1101/2024.04.03.587743

**Authors:** Yongzhi Lu, Qi Yang, Ting Ran, Guihua Zhang, Wenqi Li, Peiqi Zhou, Jielin Tang, Minxian Dai, Jinpeng Zhong, Hua Chen, Pan He, Anqi Zhou, Bao Xue, Jiayi Chen, Jiyun Zhang, Sidi Yang, Kunzhong Wu, Xinyu Wu, Miru Tang, Wei K. Zhang, Deyin Guo, Xinwen Chen, Hongming Chen, Jinsai Shang

**Affiliations:** Guangzhou National Laboratory, Guangzhou 510005, China; School of Basic Medical Sciences, Guangzhou Laboratory, Guangzhou Medical University, Guangzhou 511436, China; State Key Laboratory of Respiratory Disease, Guangzhou Medical University, Guangzhou, 511436, China; Lead contact

## Abstract

The RNA-dependent RNA polymerase (RdRp), 3C-like protease (3CL^pro^), and papain-like protease (PL^pro^) are pivotal components in the viral life cycle of SARS-CoV-2, presenting as promising therapeutic targets. Currently, all FDA-approved antiviral drugs against SARS-CoV-2 are RdRp or 3CL^pro^ inhibitors. However, the mutations causing drug resistance have been observed in RdRp and 3CL^pro^ from SARS-CoV-2, which makes it necessary to develop antivirals with novel mechanisms. Through the application of a structure-based drug design (SBDD) approach, we discovered a series of novel potent non-covalent PL^pro^ inhibitors with remarkable *in vitro* potency and *in vivo* PK properties. The co-crystal structures of PL^pro^ with leads revealed that the residues D164 and Q269 around the S2 site are critical for improving the inhibitor’s potency. The lead compound GZNL-P36 not only inhibited SARS-CoV-2 and its variants at the cellular level with EC50 ranging from 58.2 nM to 306.2 nM, but also inhibited HCoV-NL63 and HCoV-229E with EC50 of 81.6 nM and 2.66 μM, respectively. Oral administration of the compound resulted in significantly improved survival and notable reductions in lung viral loads and lesions in SARS-CoV-2 infection mouse model, consistent with RNA-seq data analysis. Our results indicate that PL^pro^ inhibitor is a promising SARS-CoV-2 therapy.

## Introduction

Over 670 million people have been infected and over 6.8 million people have died in the worldwide pandemic caused by SARS-CoV-2 virus according to data from Johns Hopkins University. Despite the efficacy demonstrated by vaccines and targeted small molecule drugs in preventing and treating COVID-19, the ongoing emergence of viral mutations, such as Alpha, Beta, Gamma, Delta and Omicron presents escalating challenges ^1–6^. The high mutation frequency of spike protein is responsible for the escape of SARS-CoV-2 from the vaccines. Unlike spike protein, the non-structural proteins (such as 3CL^pro^(nsp5) and PL^pro^(nsp3)) remain conserved among coronavirus functional proteins and show much lower mutation frequency in natural SARS-CoV- 2 variants^7,8^. The high conservativeness makes 3CL^pro^ and PL^pro^ attractive drug targets. At present, there are several clinical available small molecule anti-SARS-CoV-2 drugs. Among these drugs, remdesivir, molnupiravir, and VV116 target RNA-dependent RNA polymerase (RdRp)^9–11^, nirmatrelvir, ensitrelvir, atilotrelvir and leritrelvir target chymotrypsin-like protease (3CL^pro^, also referred as 3CL ^pro^) ^12–15^. Unfortunately, the clinical efficacy of remdesivir is controversial^16^, and multiple reported cases have already outlined an increasing observed resistance to remdesivir in immuno-compromised patients undergoing treatment with the drug^17–19^. Molnupiravir is not authorized to be used for patients under the age of 18 due to its bone and cartilage toxicity, and also not applicable for pregnant patients due to the potential risk of major birth defects and miscarriage^20^. In order to increase the half-life and the *in vivo* concentration of nirmatrelvir, ritonavir is included as a boosting agent to inhibit the activity of cytochrome P450 3A4 (CYP3A4)^21^. Ensitrelvir, the 2^nd^ generation 3CL ^pro^ inhibitor, showed favorable clinical antiviral efficacy, albeit having potent CYP3A4 inhibitory activity^13,22,23^. Although 3CL^pro^ is known as a well conserved protein, Duan *et al*. reported several potential mutant sites by which SARS-CoV- 2 might evolve the resistance to nirmatrelvir^24^.

PL^pro^, a major functional domain in SARS-CoV and SARS-CoV-2 non-structural protein 3 (nsp3), is an essential enzyme involved in viral replication and immune evasion^25–27^. PL^pro^ plays an important role in viral transcription and replication by cleaving the peptide bonds in the viral polyprotein, while the deubiquitinating and deISGylating activity of PL^pro^ is related to the immune evasion by antagonizing the host’s innate immune response upon viral infection^26^. PL^pro^-mediated deubiquitination of STING disrupted the STING-IKKε-IRF3 complex by removing the K63- linked polyubiquitin chain from LYS^289^ of STING^27^. Subsequently, the IFN-I signal pathway was inhibited. Hence, PL^pro^ is a promising drug target against SARS-CoV-2, too.

GRL0617, a SARS-CoV PL^pro^ inhibitor, also shows inhibition activity for SARS-CoV-2 PL^pro28,29^. In addition, several other SARS-CoV-2 PL^pro^ inhibitors were reported^30–35^. GRL0617 prevents the substrate binding by inducing the conformation change of Y268 on the BL2 loop that closes the BL2 loop and narrows the binding cleft^36^. However, these reported compounds only show enzymatic activity from μM to sub-μM. Except for F0213, there is no drug-like PL^pro^ inhibitors have reported *in vivo* antiviral efficacy in SARS-CoV-2 infected animal model^8,21,35^. In this study, we synthesized a series of novel PL^pro^ inhibitors and evaluated their activities. The lead compound (GZNL-P36) showed excellent *in vitro* potency as well as decent oral *in vivo* pharmacokinetic (PK) properties, more importantly, it also demonstrated similar *in vivo* antiviral efficacy in SARS-CoV-2 infected mice with ensitrelvir.

## Results

### Structure based discovery and optimization of novel PL^pro^ inhibitors

As shown in the co-crystal structure of GRL0617, the compound is bound at a shallow pocket on the protein surface (**Fig. 1A**), which has been recognized as the substrate binding site. The naphthalene ring sandwiched in the BL2 groove (*i.e.*, S1 site) is a critical group for the binding of GRL0617. In previous studies, replacing this group with alternative aromatic ring systems can somewhat maintain its bio-activity rather than increase it^34^. However, we hypothesize the potential enhancement of binding affinity through the substitution of naphthalene with a bulkier substitute, as the naphthalene ring has not yet fully occupied the entire surface of the groove, in particular the area corresponding to the residue P247. Thus, we introduced the 1,2-dihydroacenaphthylene (DHAN) group as a replacement of naphthalene ring to expand the hydrophobic contact surface with P247, and GZNL-P1 was synthesized (**Fig. 1C**). Encouragingly, its inhibitory activity (IC50 = 2.83 μM) is better than GRL0617 (IC50 = 4.82 μM) (**Fig. 1D**), indicating the beneficial effect of introducing a bulkier substitute in the BL2 groove. What’s even more interesting is that the activity can be further improved by over 10 folds for the GZNL-P3 (IC50 = 185.80 nM) when we simultaneously changed the substituent in the linking group (L) from methyl to cyclopropyl (**Fig. 1C and D**). Notably, cyclopropyl group is a much better substituent than the di-methyl as the IC50 of GZNL-P2 is just around 6.90 μM. Therefore, GZNL-P3 serves as a good starting point for further lead optimization.

**Fig. 1.**
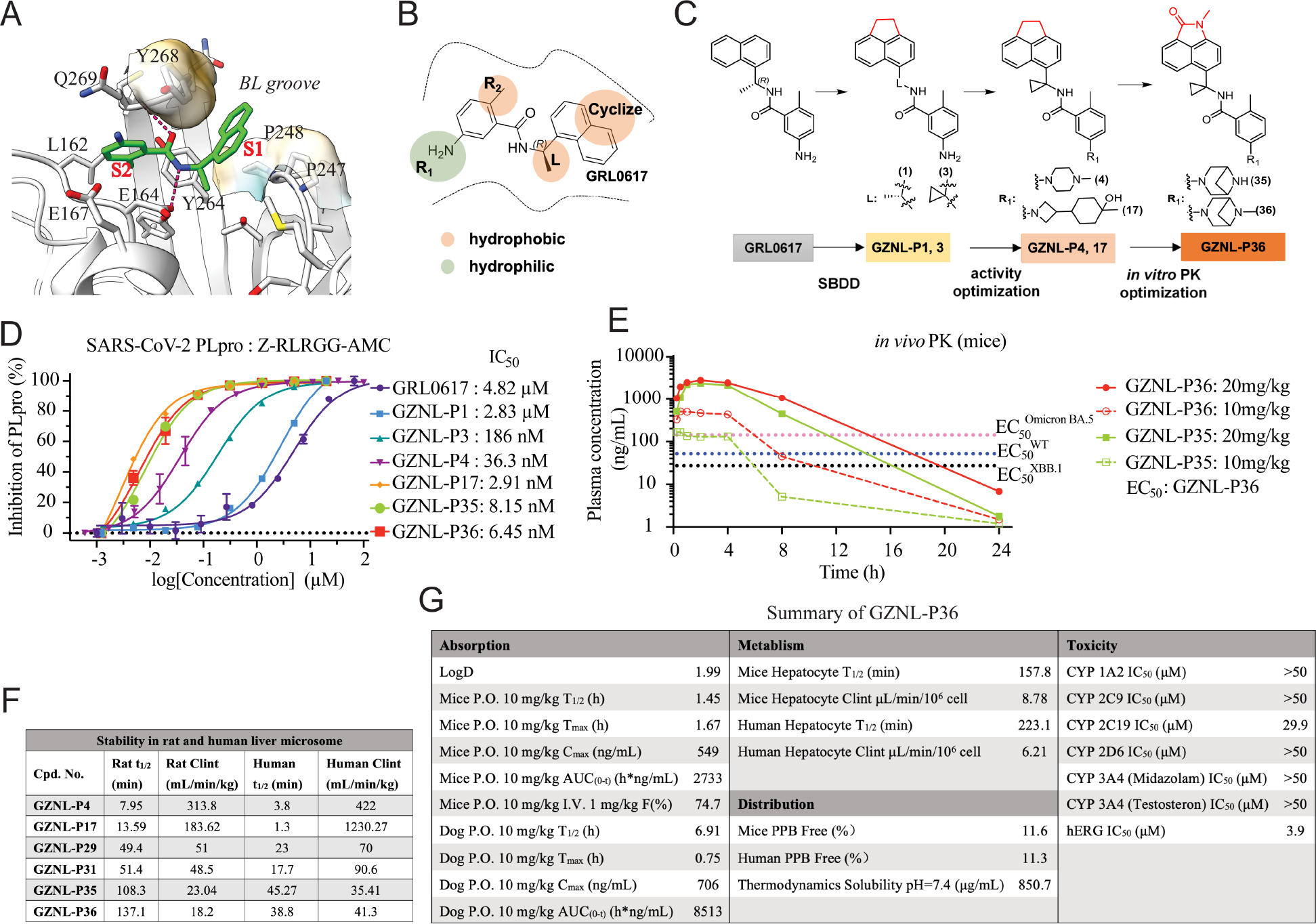
Rational design of SARS-CoV-2 PL^pro^ inhibitors and PK profiling. (**A**) Analysis of potential ligand binding sites S1 and S2 (PDB: 7JRN). The critical hydrogen bonds between GRL0617 and PL^pro^ are shown as marine dash lines. GRL0617 and the key residues of the ligand binding pocket are shown as cyan sticks and wheat sticks, respectively. (**B**) Strategies of structure- guided compound design. The optimized groups are shown as colored circles, bright orange for hydrophobicity, and split pea for hydrophilicity. (**C**) Procedures of activity optimization and *in vitro* PK optimization indicated by the representative compounds. (**D**) The inhibition activity on SARS-CoV-2 PL^pro^ of the representative compounds. (**E**) The *in vivo* PK profiling of GZNL-P35 and GZNL-P36 at 10 mg/kg and 20 mg/kg. (**F**) The *in vitro* stability in liver microsome of the representative compounds. (**G**) Summary of *in vivo* PK, metabolism, distribution, and toxicity properties of GZNL-P36.

Previous work done by Shen *et al*. has shown that it is possible to engage positively charged amine groups on the benzene of GRL0617 to interact with E167 at the S2 site^34^. This interaction could enhance activity by forming salt bridge and hydrogen bonding. A docking calculation based library design was carried out to explore R1 groups (**Fig. 1B, C**) which can form salt bridge with E167. Around 41 library compounds were selected based on docking score for synthesis and it was found that piperazine derivatives generally exhibit strong inhibition on PL^pro^). Meanwhile, compounds with 3-substituted azetidines also shows excellent activity against PL^pro^. Remarkably, GZNL-P17 is the most potent compound with IC50 of 2.91 nM, which is more potent than the best piperazine derivative, *i.e.*, GZNL-P4 (IC50 = 36.29 nM). However, modifying the methyl group on the para position of the benzene ring (*i.e.*, R2 group in **Fig. 1C**) extending to the recently identified important residue L162^37^ leads to activity drop-off. At this stage, we have successfully achieved potent enzymatic activity which is hundreds of times improved comparing to GRL0617. The most active compounds GZNL-P4 and GZNL-P17 were then selected to measure their antiviral activity against both wild-type SARS-CoV-2 and its two epidemic variants (**Extended Data** Fig. 4) in infected VeroE6 cells and sub-micromolar anti-viral potency were achieved which is much improved comparing with GRL0617 (EC50 = 23.64 μM ^30^). To evaluate their ADME properties, *in vitro* liver stability of GZNL-P4 and GZNL-P17 (**Fig. 1F**) was measured. In rat liver microsomes, the half-life time (T1/2) of both compounds is lower than 15 minutes and their intrinsic clearance rate (Clint) is very high. Moreover, they have worse stability in human liver microsomes. Metabolite identification work of GZNL-P4 indicates that compound instability could partially be attributed to the oxidation of DHAN ring (see metabolites analysis in **Extended Data** Fig. 1). To address the liver stability problem, we changed the DHAN ring to the benzoindolone ring. To our delight, for GZNL-P35 and 36, the liver stability was considerably improved (**Fig. 1F**), while their enzymatic activities were maintained at the same level. The enzymatic inhibition activities of the finally designed compound GZNL-P35 and GZNL-P36 were 8.15 nM and 6.45 nM (**Fig. 1D**), respectively. The inhibitors can stabilize and increase the melt temperature (Tm) of SARS-CoV-2 PL^pro^ (data was not shown). The cellular antiviral activity for benzoindolone compounds was examined, where GZNL-P31, 35, and 36 exhibit better potency than 100 nM against XBB.1 strains, and their toxicity to normal cells (HEK293T CC50 = 157.4, 67.67, 88.41 μM, respectively) is higher than 60 μM (Extended Data Table 1). Overall, GZNL-P35 and 36 demonstrate the most favorable profile in terms of potency and liver metabolic stability. The overall workflow for lead optimization is shown in **Fig. 1B, C**. Bioactivity data of selected compounds are listed in **Fig. 1F** and **Extended Data Table S1**.

An *in vivo* PK study was carried out for GZNL-P35 and 36 using 3 male CD1 mice (SPF level) per group with a dosage of 10 mg/kg, both compounds can reach the maximum plasma concentration at 1.58 and 1.67 h (Tmax), respectively with a peak plasma concentration (Cmax) of 227 and 549 ng/mL (**Fig. 1E**). However, the clearance of GZNL-P35 (T 1/2 = 0.96 h) is much faster than GZNL-P36 (T 1/2 = 1.45 h). This results in an enhancement of the performance of GZNL-P36 on drug blood exposure. Particularly, the bioavailability (F%) of GZNL-P36 is much higher than GZNL-P35. Further profiling of PK properties demonstrated that GZNL-P36 has weak inhibition on major metabolic enzymes in liver (**Fig. 1G**). Its inhibition on CYP 1A2, 2C9, 2C19, 2D6, 3A4 all are very weak. Additionally, its hERG toxicity is within acceptable limits. In summary, the strong *in vitro* activity and good PK properties of GZNL-P36 make it suitable to move forward to *in vivo* efficacy study.

### **X-** ray crystal structures of SARS-CoV-2 PL^pro^ with inhibitor

To clarify the binding mechanism of inhibitors, the X-ray complex crystal structures of SARS-CoV-2 PL^pro^ with GZNL-P4, GZNL-P28, GZNL-P31, and GZNL-P35 were determined (resolution range of 1.7 to 2.6 Å; **Fig. 2**, **Extended Data Table 2**). These compounds have similar binding patterns, and the unbiased electron density for PL^pro^ inhibitors GZNL-P4, GZNL-P28, GZNL-P31, and GZNL-P35 are unambiguous. The amide structures of GZNL-P4, GZNL-P28, and GZNL-P31 form two hydrogen bonds with Q269 and D164. While GZNL-P35 forms hydrogen bonds between two amide groups with D164 and E167. Compared to the naphthalene ring of GRL0617, the characteristic tricyclic group (DHAN or benzoindolone group) at the BL2 groove makes a similar π-π stacking interaction with Y268 and further expands its contact surface with P247 and P248 (**Fig. 2**). For these compounds, the substitution with methyl-substituted piperazine at R^1^ makes additional hydrogen bond and salt bridge with the residue E167 and Q269 that is responsible for the significant potency enhancement (**Fig. 2**). Based on the remarkable enhancement of enzymatic activity and the structure comparison (Extended Data Table 1) of GZNL-P1 and GZNL-P3, it is clear that the cyclopropyl group is critical for the enzymatic inhibition activity improvement. Comparison among co-crystal structures of PL^pro^ with GRL0617 (PDB: 7JRN), GZNL-P4 and GZNL-P35 shows that the cyclopropyl group makes the plane of amido bond rotate 42.7 degrees for GZNL-P4 and 34.2 degree for GZNL-P35 (**Extended Data** Fig. 2J, **2K**) comparing to that of GRL0617. This conformation change results that both GZNL- P4 and GZNL-P35 form a hydrogen bond with the residue Y264 (**Extended Data** Fig. 2D-2F). In addition, the H-π interaction between the cyclopropyl group and the aromatic side chain of Y264 is another favourable factor.

**Fig. 2.**
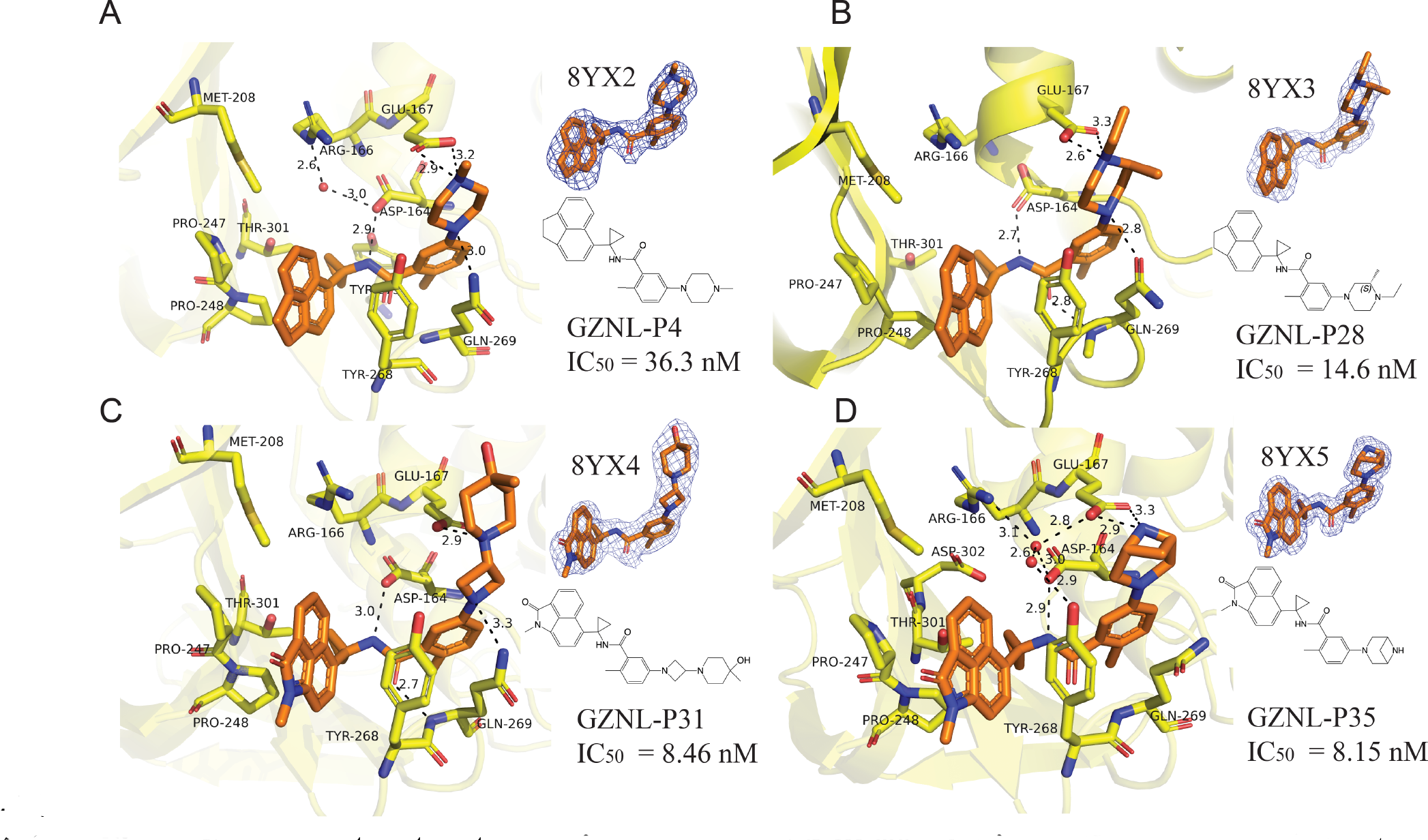
X-ray crystal structures with SARS-CoV-2 PL^pro^ inhibitors. X-ray co-crystal structure of SARS-CoV-2 PL^pro^ with GZNL-P4 (**A**), GZNL-P28 (**B**), GZNL-P31 (**C**), and GZNL-P35(**D**). The residues interacting with the ligand are shown as yellow sticks, GZNL-P4 (PDB: 8YX2), GZNL-P28 (PDB: 8YX3), GZNL-P31 (PDB: 8YX4), and GZNL-P35 (PDB: 8YX5) are shown as brown sticks. Hydrogen bonds are shown as black dashed lines and the water molecules are shown as small red spheres. The distances of hydrogen bonds and the residues are labeled.

### The mechanism of GZNL-P36 in inhibiting SARS-CoV-2 PL^pro^

It is known that PL^pro^ plays a key role in the proteolytic processing of viral polyproteins and the dysregulation of the host immune response. The deubiquitylation and de-ISGylation activity of PL^pro^ is related with the host innate immune pathways and the innate immune evasion of SARS- CoV-2^26^. To characterize the enzymatic inhibition of the designed compounds, we performed the PL^pro^ enzymatic assay using the labeled peptide substrate RLRGG-AMC (GLPBIO, GA23715). The final selected candidate compound GZNL-P36 showed potent enzymatic inhibition with IC50 value of 6.4 nM, compared to 4.8 μM for reference compound GRL0617 and 36.3 nM for the lead compound GZNL-P4 (**Fig.1D**). To investigate the thermodynamic profile of the binding between ligands and PL^pro^, we performed isothermal titration calorimetry (ITC) experiments. The measured binding affinities of GRL0617, GZNL-P35 and GZNL-P36 with PL^pro^ are 2.59 μM, 4.43 nM and 21.8nM, respectively (**Extended Data** Fig. 3A-3D). The structure optimization from GRL0617 to GZNL-P35 and GZNL-P36 is mainly driven by both enthalpy (ΔH) and entropy (−TΔS), the improvement of Gibbs free energy (ΔG) from GRL0617 to GZNL-P4/GZNL-P17 and GZNL- P35/GZNL-P36 is benefited from the substitution of naphthalene by tricyclic group of DHAN or benzoindozolone and the additional N-methyl-substituted piperazine group (GZNL-P4) or bridged piperazine (GZNL-P35) that contribute bigger interaction area with PL^pro^ (**Extended Data** Fig. 2A-2C). In addition, the piperazine group also forms hydrogen bond with residue E167 (**Fig. 2**). The increase of van der Waals interactions and the additional hydrogen bond formation are responsible for the decrease of enthalpy in the structure optimization^38^. To investigate the binding kinetic properties between ligand and receptor, biolayer interferometry (BLI) experiment was performed to get the association constant (Kon), dissociation constant (Koff) and equilibrium dissociation constant (Kd) of GRL0617, GZNL-P4, GZNL-P36 binding to PL^pro^. The Kd tested by BLI are consistent with that from ITC, with the value of 8.1 μM, 114.0 nM and 22.2 nM for GRL0617, GZNL-P4 and GZNL-P36, respectively (**Extended Data** Fig. 3E-3G). The change of Kon from GRL0617 to GZNL-P4 and GZNL-P36 are contributed by the increasing hydrophobic force due to the substitution of naphthalene by a bigger tricyclic acenaphthylene group and the additional piperazine group, by the increasing polar contact due to the formation of hydrogen bond between the piperazine group of GZNL-P4/GZNL-P36 and the residue E167^38^. The melt temperature (Tm) of PL^pro^ determined by differential scanning fluorimetry (DSF) assay indicated that PL^pro^ was significantly stabilized by the compound binding, the ΔTm of PL^pro^ with or without incubation of GRL0617, GZNL-P4, GZNL-P19, GZNL-35, and GZNL-P36 increased by 16 to 22.5 ℃ (data was not shown). The enzymatic inhibition activity is related with the blockade of substrate binding to PL^pro^ by inhibitors. GRL0617 bound to PL^pro^ mainly hinders the residue L73 of substrate (72RLRGG76), but for GZNL-P4 and GZNL-P35, piperazine group also impedes the residue R72 of the substrate (**Extended Data** Fig. 2G-2I).

### Evaluation of *in vitro* antiviral activity cross coronavirus family

To test whether GZNL-P36 could effectively inhibit PL^pro^ across multiple coronavirus subtypes, we performed a fluorescence resonance energy transfer (FRET) inhibition assay against PL^pro^ proteins from different species coronaviruses from genera alpha-, beta-, gamma-, and deltacoronaviruses (**Fig. 3A,B**). GZNL-P36 exhibited broad maximum inhibition efficacy against PL^pro^ derived from all coronaviruses tested (**Fig. 3A**). The cellular antiviral activity of GZNL-P36 was examined by a cell protection assay. In this assay, the cytopathic effect (CPE) of SARS-CoV- 2-infected Vero E6 cells with or without treatment by the compounds was assessed using Celigo Image Cytometer^39^. The cells were challenged with WT SARS-CoV-2 and two other variants Omicron BA.5 and XBB.1. GZNL-P36 dose-dependently protected cells from death with 50% effective concentration (EC50) values for wild type (WT), Omicron BA.5 and XBB.1 is 111.0 nM, 306.2 nM and 58.6 nM, respectively (**Fig. 3C-3E**). Compared with S-217622 (Ensitrelvir), GZNL- P36 possessed similar effective anti-viral activity for SARS-CoV-2 and its variants (**Fig. 3C-3E**). Besides SARS-CoV-2, GZNL-P36 also illustrated anti-viral activity for the other coronaviruses, such as HCoV-NL63 (EC50: 81.6 nM), HCoV-229E (EC50: 2.66 μM) and HCoV-OC43 (EC50: 46.3 μM) (**Fig. 3F**). Together, our data demonstrated that GZNL-P36 rendered superb cross-protection against SARS-CoV-2, HCoV-OC43, HCoV-229E, and HCoV-NL63, exhibiting potent broad-spectrum anticoronaviral efficacy.

**Fig. 3.**
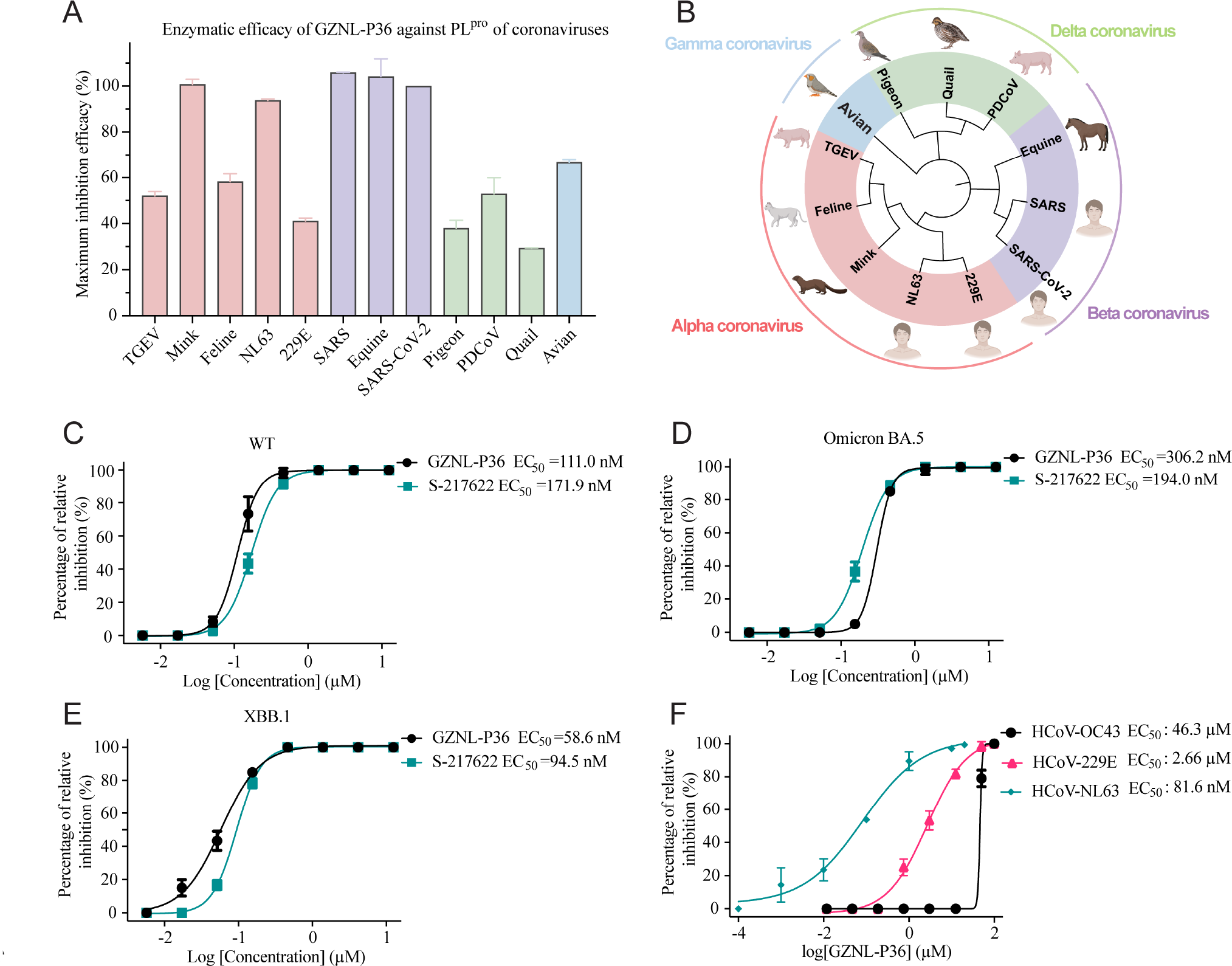
Pan-antiviral activity of GZNL-P36 against SARS-CoV-2 variants and other coronavirus. (**A**) Enzymatic maximum inhibition efficacy of GZNL-P36 against coronaviruses PL^pro^. (**B**) The phylogenetic tree of coronaviruses PL^pro^ used in this experiment. Antiviral activity of GZNL-P36 against SARS-CoV-2 wild type (WT) (**C**), variants Omicron BA.5 (**D**), and XBB.1 (**E**). Vero E6 cells were pre-treated with indicated compounds with different concentrations for 1 h and then infected with SARS-CoV-2 wild type (WT), variants Omicron BA.5, and XBB.1 at an MOI of 0.01. The EC50 was assessed after being cultured for three days. (**F**) The representative inhibition curves of GZNL-P36 against HCoV-229E and HCoV-OC43 in Huh-7 cells. The EC50 was assessed after being cultured for two days. Three independent experiments were performed on infections and one representative is shown.

### *In vivo* antiviral efficacy of GZNL-P36

To assess the *in vivo* anti-viral activity of GZNL-P36, we treated the model mice infected with SARS-CoV-2 XBB.1 by oral administration (**Fig. 4A**). K18-hACE2 transgenic mice aged 8 weeks were used as our mouse model, forty eight female hACE2 transgenic mice were divided into six groups with eight mice in each group to evaluate the efficacy of mock, vehicle, positive comparator S-217622 of 25 milligrams per kilograms (mpk), and GZNL-P36 of 25 mpk, 50 mpk and 100 mpk in the therapeutic treatment. The weight loss plot shows the about 15% loss of the vehicle group, but the weight loss is less than 10% for the treated groups (**Fig. 4B**). The lung live viral titers cannot be detected (**Fig. 4C**) for the group treated with GZNL-P36 at 100 mpk. The groups treated by GZNL-P36 at the dose of 25 mpk or 50 mpk also showed significant viral titer decrease at 2 days post-infection. The anti-viral efficiency of GZNL-P36 is slightly weaker than the same dose of positive drug S-217622. Immunohistochemistry assays with SARS-CoV-2 nucleocaspid protein antibody revealed that abundant expression of viral antigen was identified in the lung of vehicle-treated mice at 4 days post-infection (**Fig. 4D**). In contrast, GZNL-P36 treatment, even when administered after the virus challenge, markedly suppressed viral nucleocapsid protein expression in the lung (**Fig. 4D**). Next, haematoxylin and eosin (H&E) stained lungs indicate that the lung was significantly protected from the GZNL-P36 treatment (**Fig. 4E**).

**Fig. 4.**
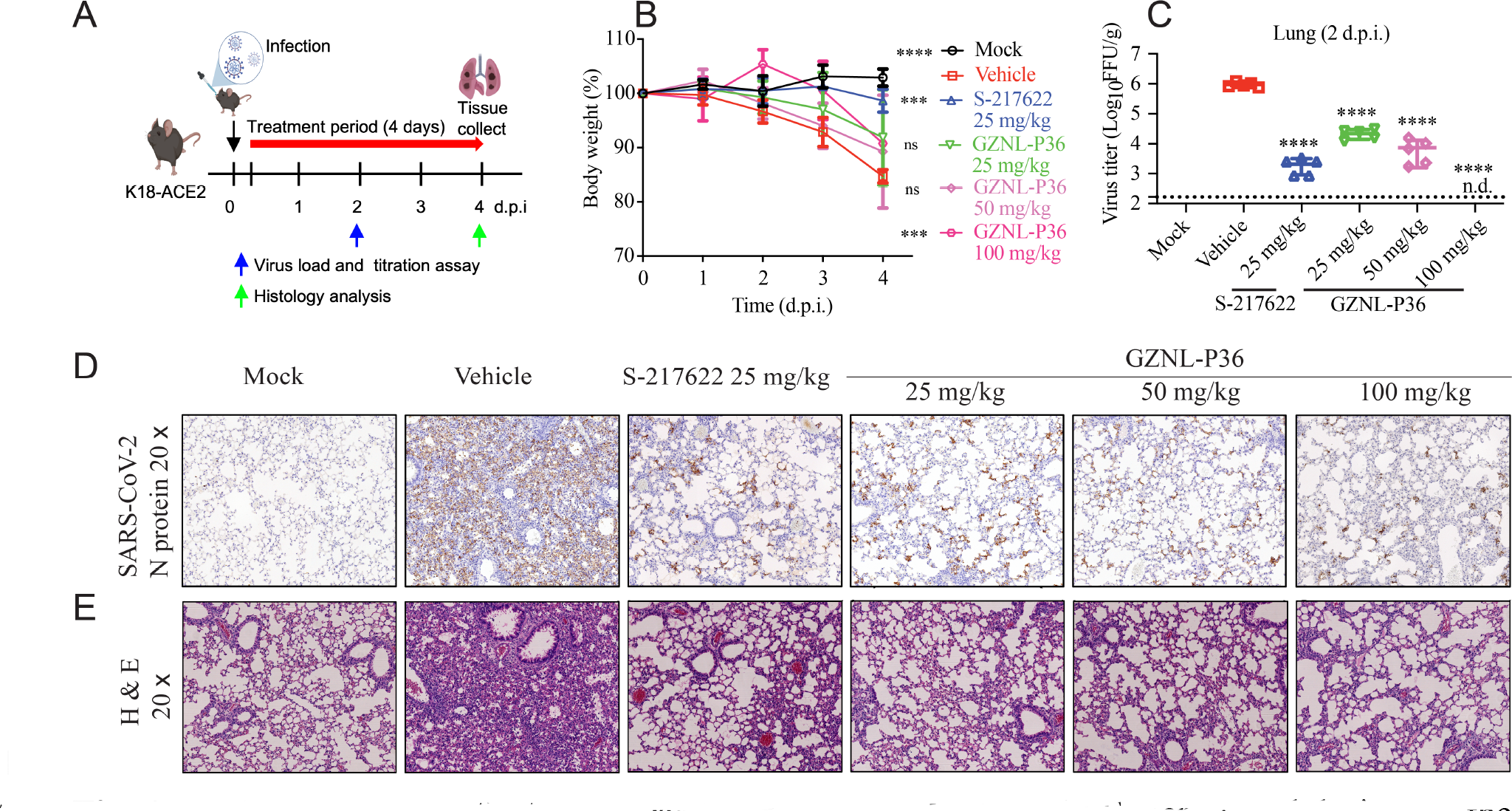
*In vivo* antiviral activity of PL^pro^ inhibitor GZNL-P36. (**A**) Experimental design for the 4-day experiment in K18-ACE2 mouse. (**B**) Body weight loss of mice from different groups. (**C**) Live viral titers in lungs collected at 2 d.p.i. (**D**) and (**E**) Lungs collected at 4 d.p.i. from different groups were immunostained with SARS-CoV-2 nucleocaspid protein antibody (**D**) or stained with haematoxylin and eosin (H&E) (**E**). Each dot represents one mouse at the indicated time point. The data are representative of at least two experiments. The error bars are mean ± SD. Statistical differences were determined by two-way ANOVA in **C**. \*\*\**P* < 0.001, \*\*\*\**P* < 0.0001; ns, not significant.

PL^pro^ can dysregulate the host inflammation and antiviral response due to its deubiquitinating activity. The transcription levels of inflammatory genes, including CXCL10, IFNB1, and IFNγ1, were determined with the lungs of the mice collected at 2 d.p.i. Compared to the vehicle group, the transcription level of these pro-inflammatories in GZNL-P36 treated groups was significantly decreased. Furthermore, the transcription level of CXCL10 and IFNγ1 in the GZNL-P36 treated groups was also lower than that in the S-217622 treated group (**Extended Data** Fig. 6). These results indicated that PL^pro^ inhibitors may provide more benefits on the anti-inflammation properties than S-217622.

We further perform bulk RNA sequencing on lung samples of all SARS-CoV-2 infected mice. GSVA analysis results revealed that GZNL-P36 successfully reversed most of SARS-CoV-2- induced changes, identical to the 3CL ^pro^ inhibitor S-217622 (**Fig. 5**). More importantly, high dose of GZNL-P36 greatly reversed the GSVA scores of both WP_FOXP3_IN_COVID19 and WP_PATHOGENESIS_OF SARSCOV2_MEDIATED_BY_NSPINSP10_COMPLEX genesets, while S-217622 failed, suggesting our GZNL-P36 might possess synergistic effect on the recovery of SARS-CoV-2 infected mice comparing with the 3CL ^pro^ inhibitor S-217622 (**Extended Data** Fig. 7).

**Fig. 5.**
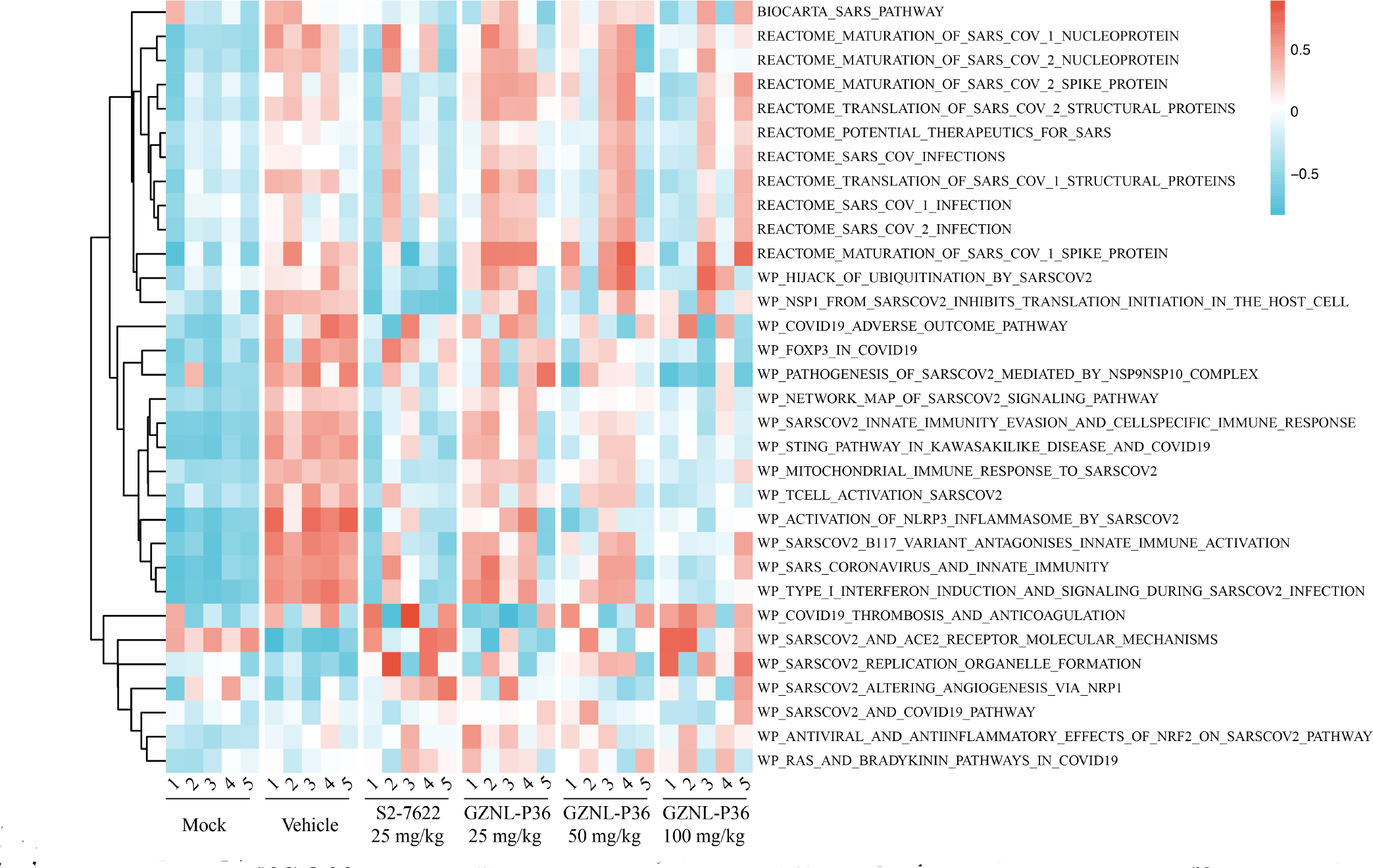
RNAseq analysis of GZNL-P36 in SARS-CoV-2 infected Mice. GSVA scores of selected genesets on bulk RNAseq data of GZNL-P36 in SARS-CoV-2 infected Mice. Genesets were obtained from MSigDB and curated based on a criterion of including keywords such as “SARS” or “COVID”. The data were then scaled, GSVA scores ranging from −1 (blue, down-regulated) to 1 (red, up-regulated).

## Discussion

Vaccines and neutralizing antibodies cannot provide complete protection against the continuously emerging variants of SARS-CoV-2. Small molecule antiviral drugs targeting the conserved viral proteases are particularly important. At present, there are several clinically available drugs targeting RdRp and 3CL^pro^. Unfortunately, the resistance mutations of RdRp^18,19^ and 3CL ^pro24,40–42^ have been reported. For PL^pro^, another important potential antiviral drug target, still no targeted drug has been reported. Starting from a weak PL^pro^ inhibitor GRL0617, a novel benzoindolone series of PL^pro^ inhibitors was discovered through the utilization of docking-based library design. Our lead compound GZNL-P36 shows excellent PL^pro^ inhibitory potency and decent pharmacokinetics and *in vitro* safety profile. Furthermore, this compound shows strong *in vivo* anti-viral efficacy in antiviral mice model suggesting its potential as a COVID-19 therapy candidate. Interestingly, compared to S-217622, GZNL-P36 treatment showed lower expression of the pro-inflammatory genes. Our results demonstrate that PL^pro^ is an attractive druggable antiviral target and PL^pro^ inhibitor is a class of promising antiviral drug with dual-effect on antiviral and anti-inflammation.

## Methods and Materials

Detailed descriptions of the in vitro pharmacology studies, *in vivo* pharmacology studies, transcriptomics studies, X-ray crystallography, computational study, and synthetic methods can be found in supplementary materials.

## Acknowledgements

We are grateful for support from Shanghai Synchrotron Radiation Facility (SSRF) beamlines (BL19U1 and BL02U1). This study was supported by grants from the National Natural Science Foundation of China (821704730), the startup and R&D Program of Guangzhou National Laboratory (YW-YWYM0202, GZNL2023A02012, GZNL2023A01008, SRPG22-002, SRPG22-003), and the Guangdong Natural Science Foundation (2021QN020451, 2021CX020227), and the Basic and Applied Basic Research Projects of Guangzhou Basic Research Program (202201011795, 2023A04J0161).

## Author contributions

Y.L., Q.Y., W.L., M.D., J.T., J.C., J.Z., A.Z., S.Y., K.W., and X.W. performed the cellular and biochemical assays. Y.L. and J.S. performed crystallography. G.Z., P.Z., H.C., P.H., and T.R. performed the medicinal chemistry. Q.Y., W.L., B.X., and J.T. performed the animal models. T.R., M.T., and C.H. performed the computational studies. Y.L., W.L., M.D., J.C., J.Z., K.W., and X.W. performed cloning and/or purified proteins. Y.L., W.L., J.S., and W.K.Z. performed the transcriptome study. Y.L., Q.Y., T.R. X.C., C.H., and J.S. conceived the experiments and wrote the manuscript with input from all authors.

## Competing interests

The authors declare no competing interests.

---

**Extended Data Table S1.**
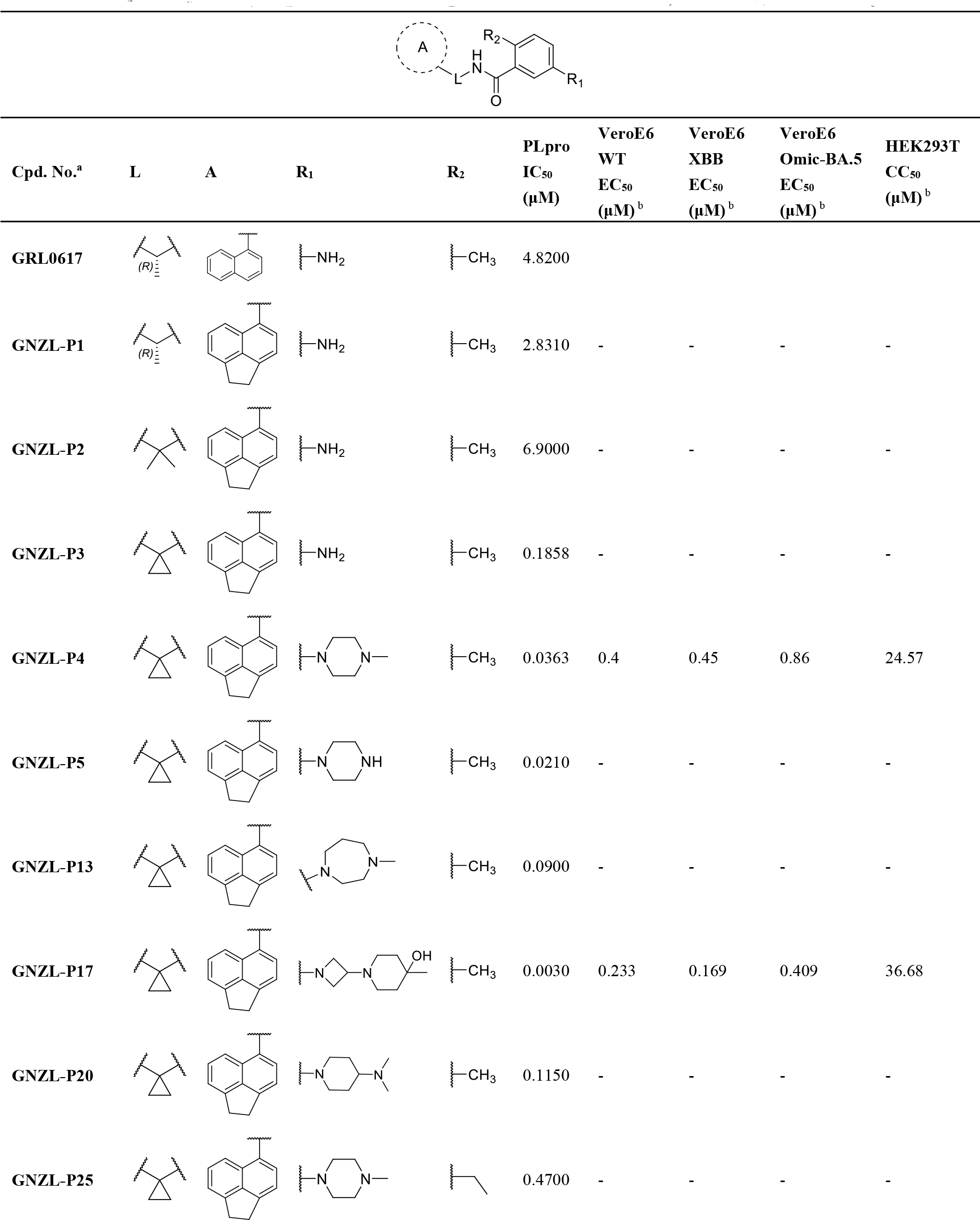
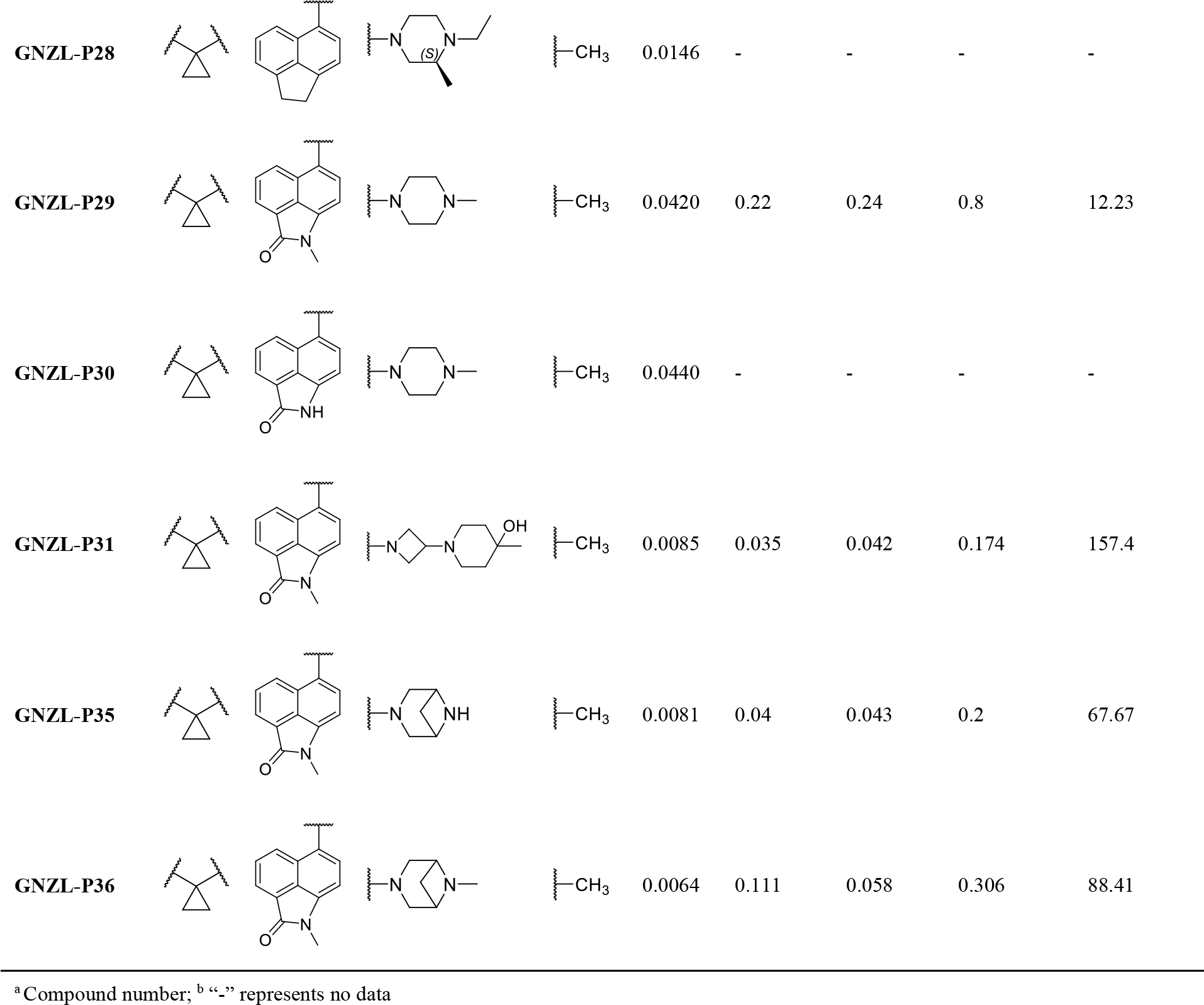
Representative compounds with enzymatic, antiviral, cell toxicity activity.

**Extended Data Table S2.**
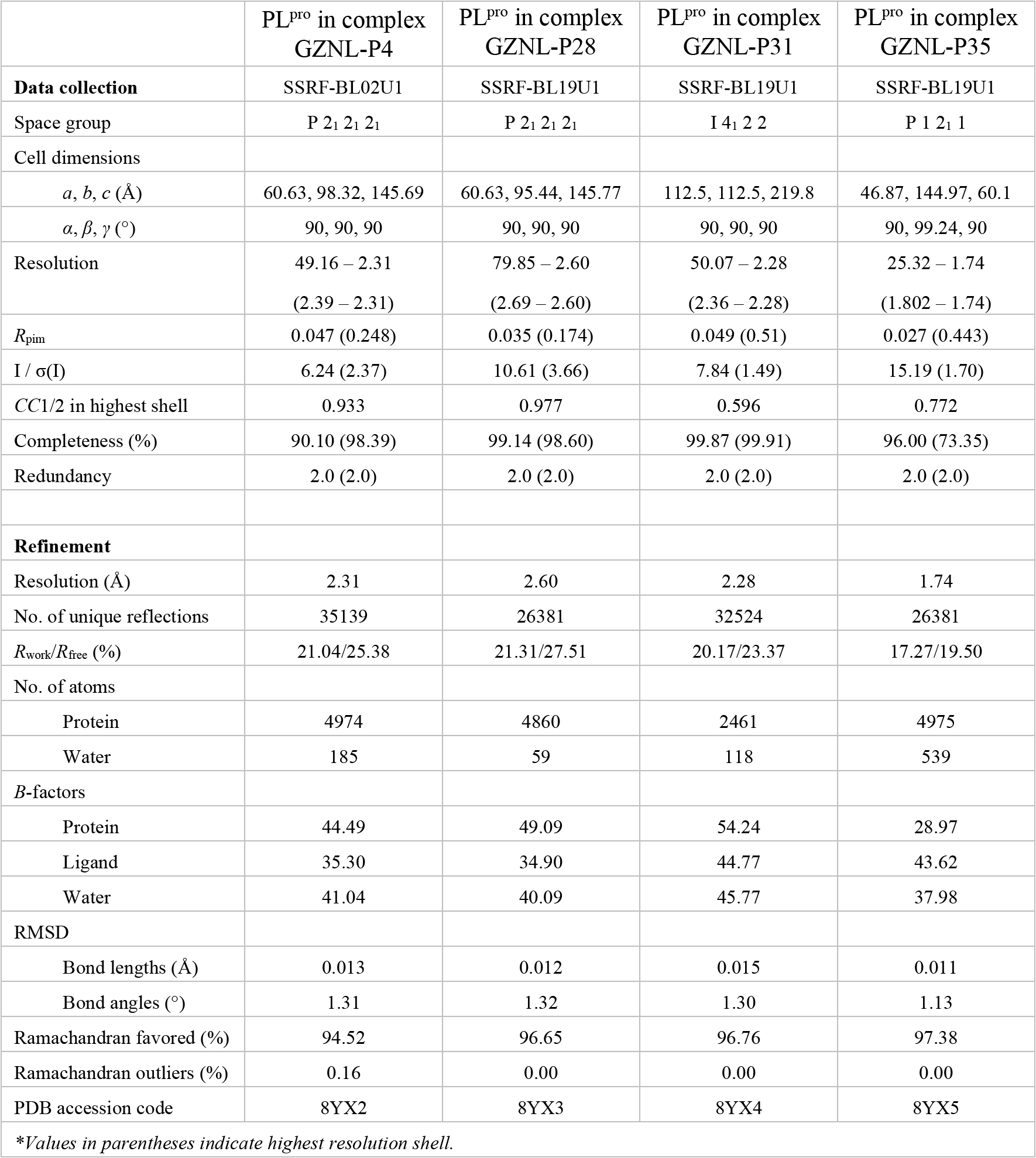
X-ray data collection and refinement statistics.

**Extended Data Fig. 1.**
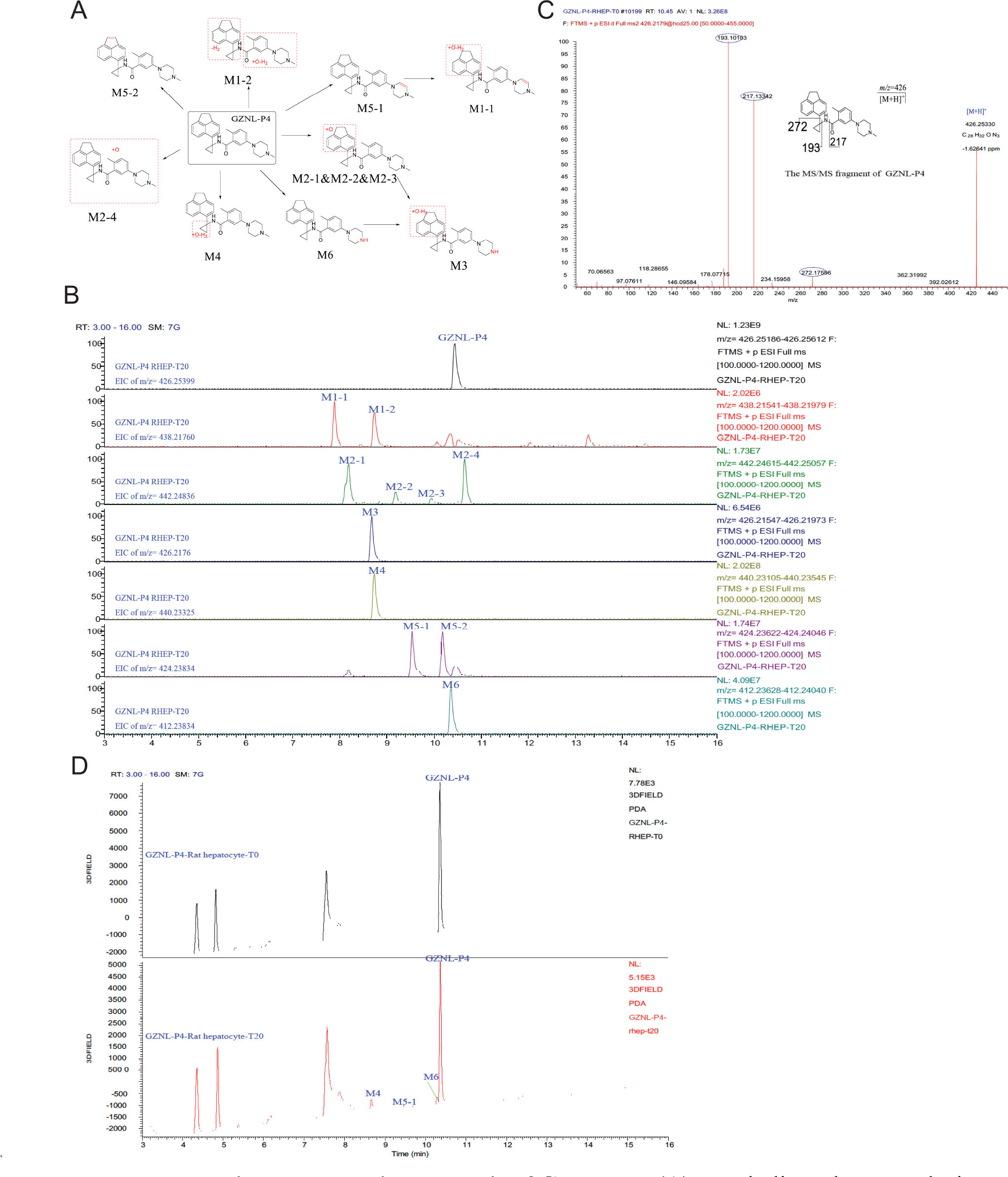
Metabolism analysis of GZNL-P4. (**A**) Metabolic pathway analysis. Sites of Metabolism are highlighted by red color. (**B**) MS detection of metabolites by retention time. (**C**) MS characterization of metabolites by mass-over-charge ratio (m/z). (**D**) UV spectrum detection of metabolites.

**Extended Data Fig. 2.**
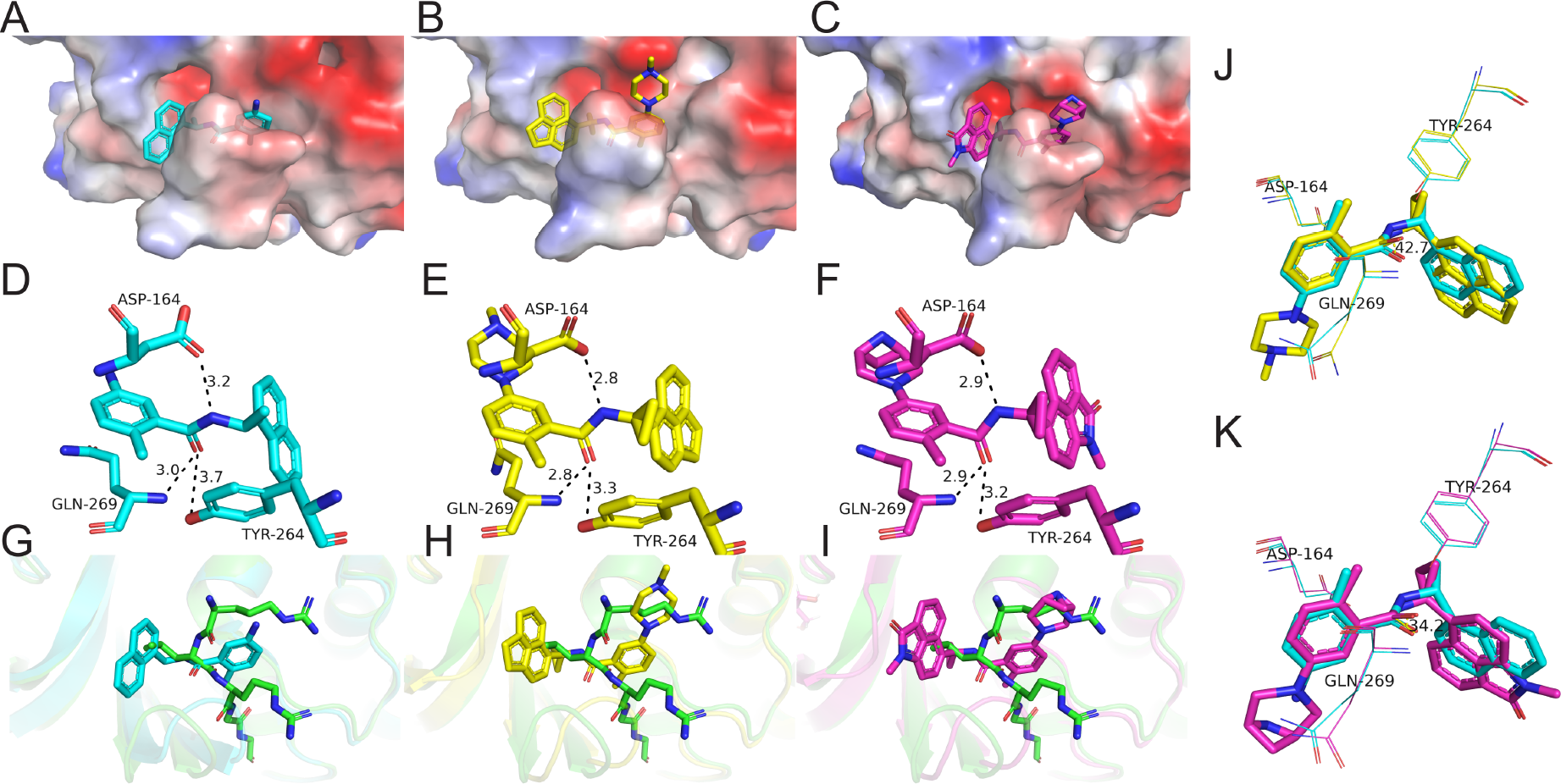
The comparison of binding conformations between GRL0617 and GZNL-P4 or GZNL-P35. (**A**) **-** (**C**) The pattern of GRL0617 (PDB: 7JRN) (**A**), GZNL-P4 (PDB: 8YX2) (**B**), and GZNL-P35 (PDB: 8YX5) (**C**) binding to SARS-CoV-2 PL^pro^. PLpro is shown as electrostatics surface and the inhibitors (GRL0617, GZNL-P4, GZNL-P35) are shown as sticks. (**D**) **-** (**F**) The comparison of GRL0617 (**D**), GZNL-P4 (**E**), and GZNL-P35 (**F**) binding to SARS-CoV-2 PL^pro^. (**G**) **-** (**I**) Comparison of the PL^pro^ substrate and GRL0617 (**G**), GZNL-P4 (**H**), and GZNL-P35 (**I**) binding to SARS-CoV-2 PL^pro^. The structures are superimposed using the PL^pro^ protein structure of the co-crystal structures. The substrate peptide RLRGG (PDB: 6YVA), GRL0617 (PDB: 7JRN), GZNL-P4, and GZNL-P35 are shown as green, cyan, yellow, and purple sticks, respectively. The protein is shown as cartoon with the same color as the ligand. (**J**) **-** (**K**) Comparison of the binding conformation between GRL0617 and GZNL-P4 (**J**) or GZNL-P35 (**K**). Hydrogen bonds are shown as black dashed lines. The distances of hydrogen bonds and the residues are labeled.

**Extended Data Fig. 3.**
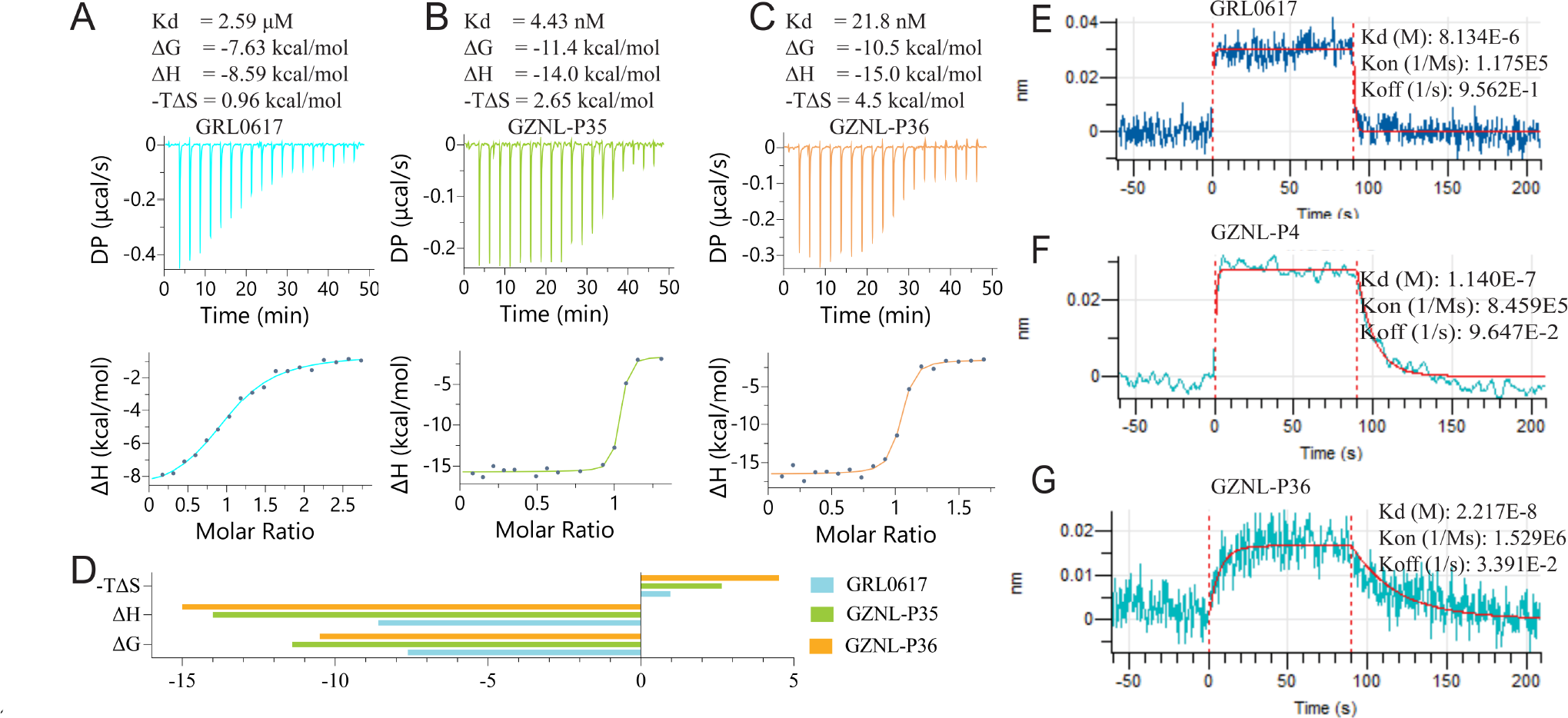
The binding constant determination by ITC and BLI. (**A**) **-** (**C**) ITC binding curve for the interactions of SARS-CoV-2 PL^pro^ with GRL0617 (**A**), GZNL-P35 (**B**), and GZNL-P36 (**C**). (**D**) The △G, △H, and -T△H for (**A**) **-** (**C**). (**E**) **-** (**G**) BLI binding curve for the interactions of SARS-CoV-2 PL^pro^ with GRL0617 (**E**), GZNL-P4 (**F**) and GZNL-P36 (**G**).

**Extended Data Fig. 4.**
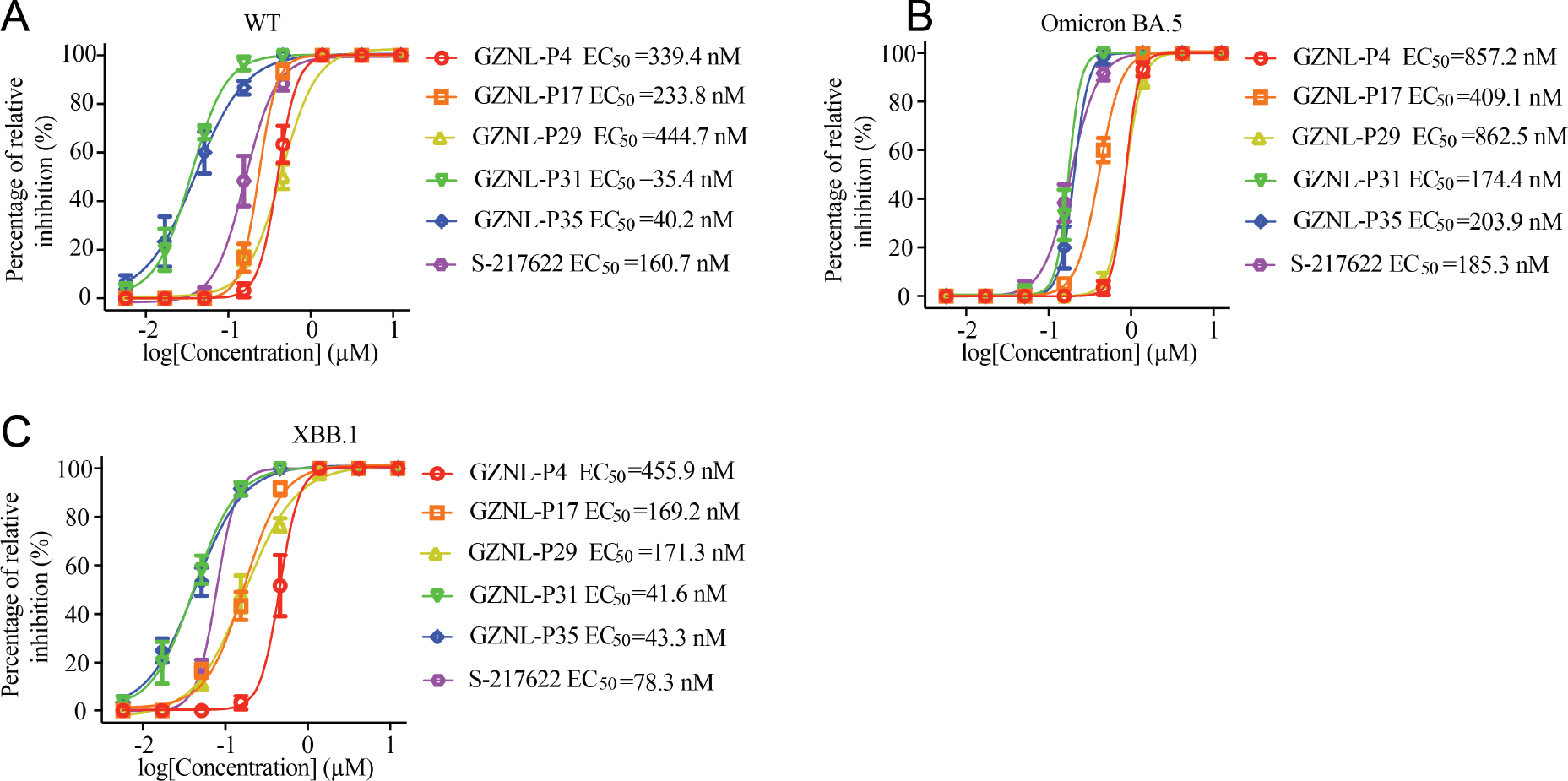
The anti-viral activity of the designed compounds against SARS-CoV-2. Vero E6 cells were pre-treated with indicated compounds with different concentrations for 1 h and then infected with SARS-CoV-2 WT (**A**), variants Omicron BA.5 (**B**), and XBB.1 (**C**) at an MOI of 0.01. The Y-axis of the graphs represents the mean inhibition (%) of virus yield of the compounds. The EC50 was assessed after being cultured for three days. Three independent experiments were performed on infections and one representative is shown.

**Extended Data Fig. 5.**
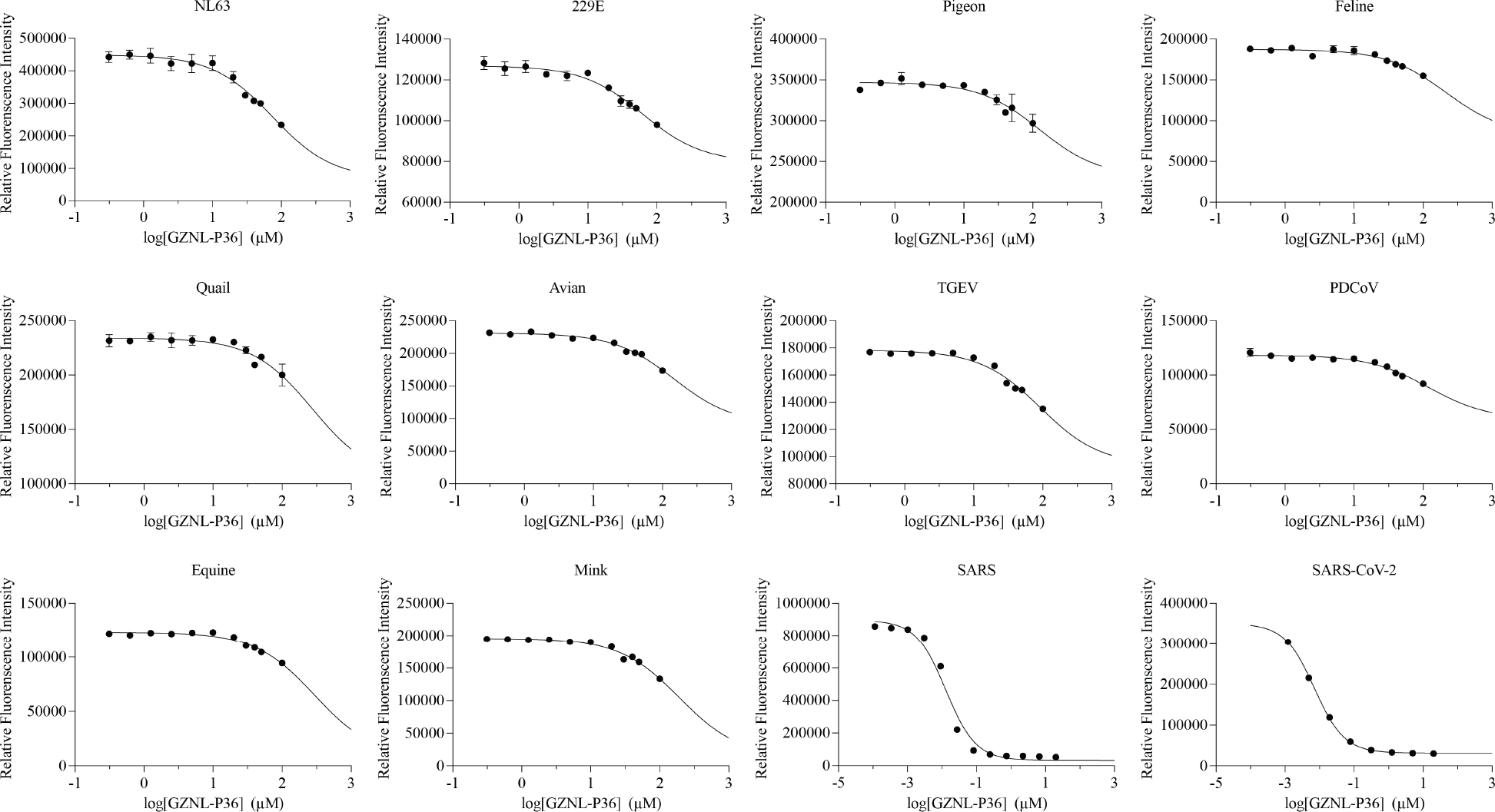
Evaluation of enzymatic inhibition efficacy of GZNL-P36 against PL^pro^ from different species coronaviruses by a fluorescence resonance energy transfer (FRET) inhibition assay. Different species coronaviruses PL^pro^ tested in this experiment are from alpha-, beta-, delta-, and gamma-coronavirus sub-family.

**Extended Data Fig. 6.**
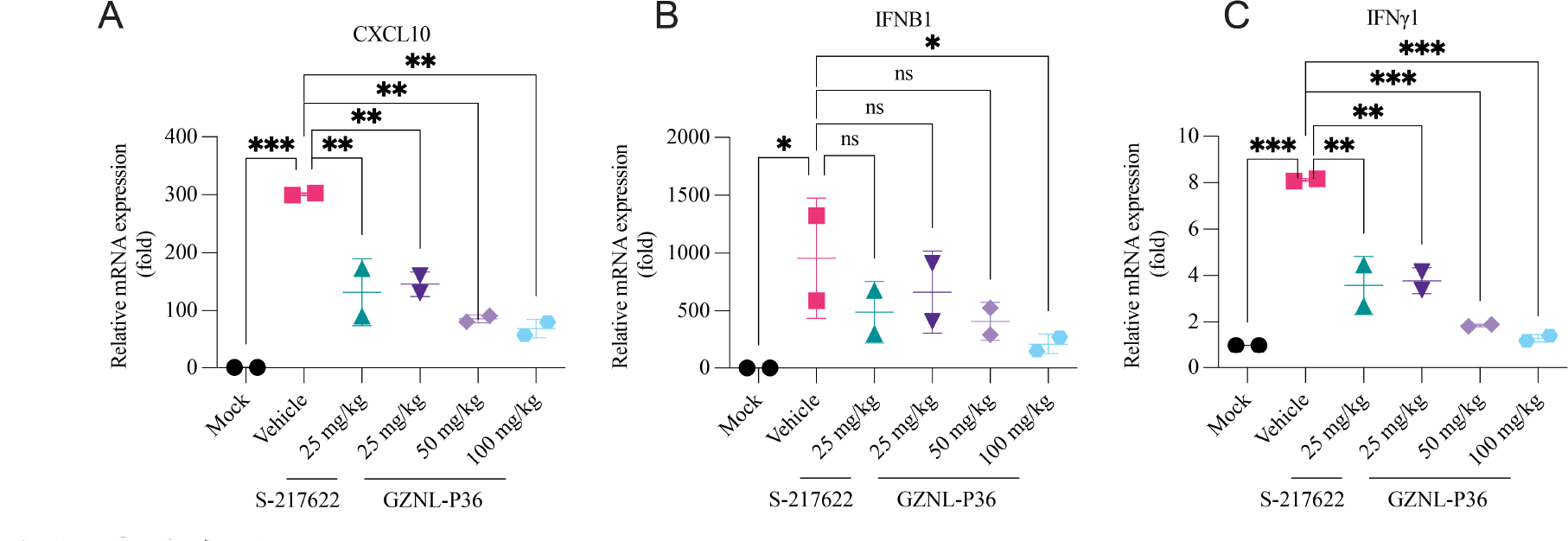
The effects of GZNL-P36 on the transcription level of anti-inflammatory genes in SARS-CoV-2 infected mice. Relative mRNA expression of CXCL10 (**A**), IFNB1 (**B**), and IFNγ1 (**C**) of the lungs collected at 2 d.p.i.. Each dot represents one mouse at the indicated time point. The error bars are mean ± SD. Statistical differences were determined by two-way ANOVA. \*\**P* < 0.01, \*\*\**P* < 0.001; ns, not significant.

**Extended Data Fig. 7.**
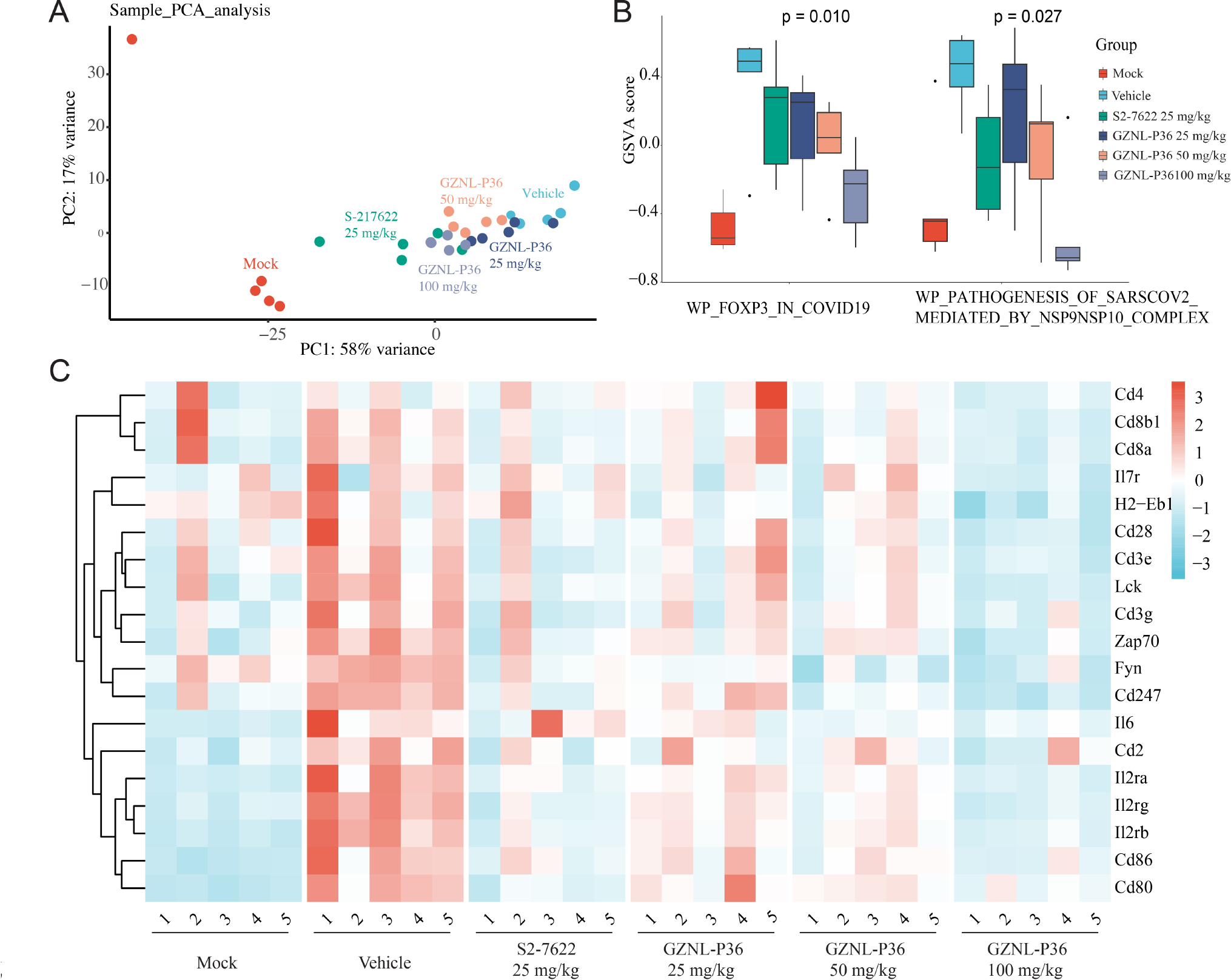
RNAseq results of GZNL-P36 in SARS-CoV-2 infected mice. Principal component analysis (PCA) of all datasets on count matrix. (**B**) GSVA scores of different treatment groups revealed a better efficacy of GZNL-P36 on these specific genesets comparing with the positive control S-217622. P value represented significance among GSVA scores of indicated datasets in the same geneset. (**C**) Relative expression levels of genes included in genesets demonstrated in (B). Scale, row-scaled TPM of certain gene (row) in certain sample (column).

## Notes

### Competing Interest Statement

The authors have declared no competing interest.

### Summary of Updates

Updated Figure legends and a few typos on the manuscript and supporting information.

