## Supporting Information for "Discovery of orally bioavailable SARS-CoV-2 papain-like protease inhibitor as a potential treatment for COVID-19"

Yongzhi Lu *et al.*

### Methods

#### Docking protocol

To explore optimum groups (*i.e.*, in $R_1$ in Figure 1C) that interact with E167 based on the DHAN scaffold of GZNL-P3, by using commercially available reagents, we designed a virtual library comprising totally 2347 compounds containing positively charged amine groups (Scheme S1). Molecular docking was then employed to predict the binding poses of the compounds. In the end, 41 top ranked compounds (by Glide docking score) that can not only maintain a similar binding mode to GZNL-P3, but also form salt bridge interaction with GLU167 in an appropriate distance (the cutoff is set to 4.0 Å) were selected for synthesis and their enzymatic activities were measured. The details of molecular docking, synthesis, and biological assays were described in the following sections.

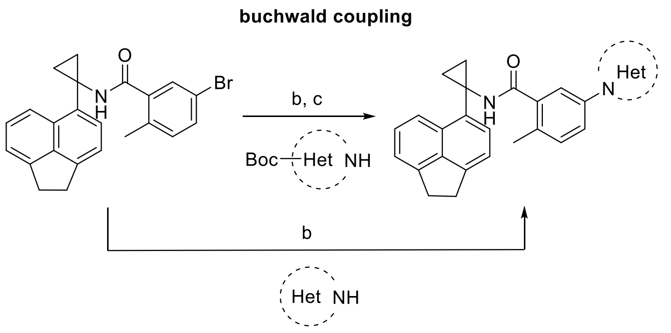


(b) Pd(OAc)2, Ru-Phos, Cs2CO3, 1,4-dioxane, 110 oC; (c) TFA, DCM, 25 oC.

**Scheme S1. Scheme for library synthesis at $R^1$ through buchwald coupling reaction.**

#### Docking method

Molecular docking was carried out using the Glide docking algorithm integrated in the software of Schrödinger (version 2020). The co-crystal structure of SARS-CoV-2 $PL^{pro}$ with GRL0617 (PDB code: 7JRN) was selected as the receptor model for docking. The protein structure was preprocessed using the Protein Preparation Wizard in the software. Bond orders were assigned to all bonds in the structure properly. Amino acids with missing side chains were also repaired. In addition, all crystal waters were removed. The Epik algorithm was chosen to generate ionization and tautomeric states at the pH of 7.0, and the structure was minimized using the OPLS3 force field until that heavy atoms were converged to 0.3 Å root mean square deviation to the original coordinates. After this, the minimized structure was then used to prepare the receptor grid for Glide docking, which defined a box space centered on the gravity of the crystal ligand as the docking site. On the other hand, ligands for docking were prepared using the LigPrep module in Schrödinger. Similar to protein preprocessing, the ligands were protonated using the Epik algorithm. Tautomers and stereoisomers were also enumerated. One lowest energy conformation per compound was generated using the OPLS3 force field. The prepared ligands were then docked to the predefined docking site. Standard docking precision was employed, where flexible mode was adopted for ligand conformation sampling. At most 10 poses per ligand were eventually reserved after docking and they were ranked by the Glidescore, which is a scoring function for pose measurement. The poses which did not map to the binding mode of GZNL-P3, including the π-π interaction within the BL2 groove and the two hydrogen bonds with Q269 and E162 mediated by the amide group were removed and the rest went through further analysis.

**Chemical synthesis**

The compounds with the scaffold of 1,2-dihydroacenaphthylene (DHAN) were synthesized following the method in Scheme S2. In brief, the synthesis started from the agent S1, which was used to synthesize the intermediate S2 with a cyano group to substitute the bromide in S1. S2 was then converted to into the key intermediate S3-3 with cyclopropylamine by Kulinkovich-Szymoniak or S3-2 with di-methylamine by grignard reagents. On the other hand, S1 was converted to S1-1 through nucleophilic addition reaction, which was then utilized to prepare the key intermediate S3-1. S3-1, S3-2 and S3-3 were respectively condensed with 5-amino-2-methylbenzoic acid to generate the products GZNL-P1-3. In addition, S3-3 reacts with the intermediate S6 by amino condensation to produce the final products GZNL-P4-5, 13, 17, 20, 25, 28 while the synthesis of the intermediate S6 follows the method in Scheme S3, which used bromo-2-methylbenzoate as raw material to synthesize the intermediate S6 by two reaction steps. At first, the bromide in the raw material S4 was substituted with a series of amine groups by Buchwald-Hartwig coupling or Suzuki reactions. Then, the intermediate product S5 at this step was hydrolyzed to generate S6. To be noted, the synthesis of GZNL-P28 follows the same method but uses different raw material, which is 1-bromo-4,5-dimethylnaphthalene.

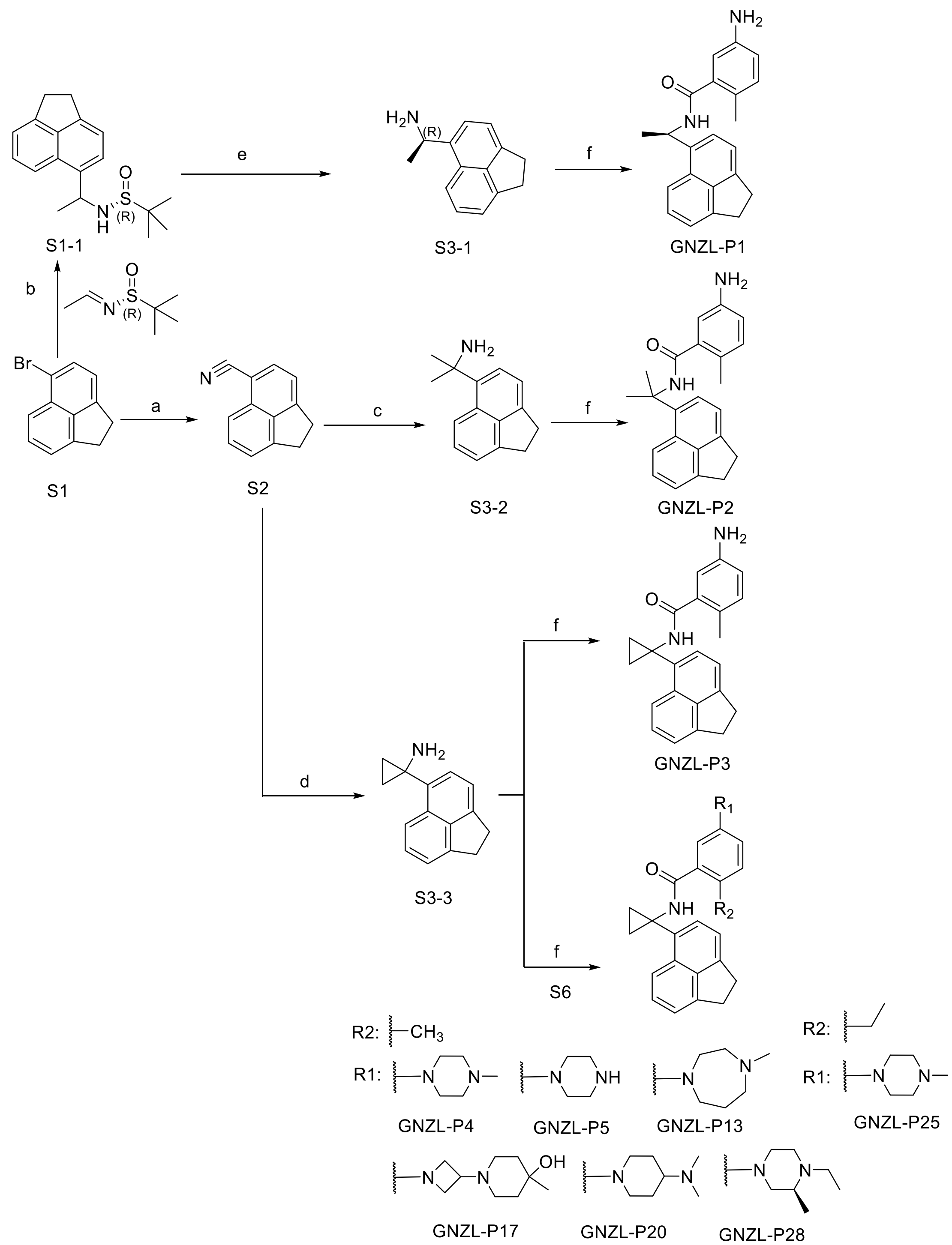


**Scheme S2. General procedure for the synthesis of compounds with the DHAN scaffold.** Reagents and conditions: **a)** CuCN, NMP, 170 ºC, 81%. **b)** n-BuLi, THF, -78 ºC-r.t., 61%. **c)** $CeCl_3$, $CH_3Li$, -78 ºC-r.t., 64%. **d)** i) Titanium isopropoxide, $CH_3CH_2MgBr$, THF, -78 ºC; ii) Triisopropyl borate, r.t., 47%; **e)** HCl/dioxane, MeOH, r.t., 100%. **f)** HATU, DIPEA, DMF, 50 ºC, 5%-80%.

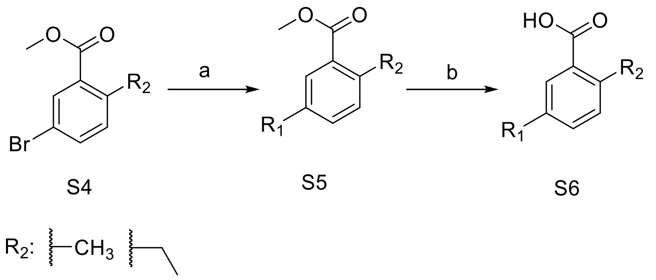


**Scheme S3. General procedure for the synthesis of chemical intermediate S6.** Reagents and conditions: **a)** $Pd_2(dba)_3$, Xphos, $Cs_2CO_3$, Dioxane, 110°C, 80%-90%. **b)** NaOH, THF/EtOH/$H_2O$, 60°C, 70%-95%.

**(R)-5-amino-N-(1-(1,2-dihydroacenaphthylen-5-yl)ethyl)-2-methylbenzamide (GNZL-P1)**. White solid, 50% yield. $^1$H NMR (600 MHz, DMSO-d6) δ 8.66 (d, J = 8.2 Hz, 1H), 7.88 (d, J = 8.4 Hz, 1H), 7.52-7.49 (m, 2H), 7.33 (d, J = 6.8 Hz, 1H), 7.28 (d, J = 7.1 Hz, 1H), 6.83 (d, J = 8.1 Hz, 1H), 6.52 (d, J = 2.4 Hz, 1H), 6.50 (dd, J = 8.0, 2.4 Hz, 1H), 5.77 (p, J = 7.1 Hz, 1H), 4.98 (s, 2H), 3.38-3.35 (m, 2H), 3.33-3.30 (m, 2H), 2.08 (s, 3H), 1.53 (d, J = 7.0 Hz, 3H). $^{13}$C NMR (151 MHz, DMSO-d6) δ 169.16, 146.59, 146.52, 144.97, 139.38, 138.28, 136.80, 131.13, 129.50, 128.25, 124.50, 121.75, 119.60, 119.41, 119.31, 115.06, 113.08, 44.37, 29.80, 22.00, 18.76. LC-MS (ESI, m/z): $C_{22}H_{22}N_2O$, $[M+H]^+$ = 331.21.

**5-amino-N-(2-(1,2-dihydroacenaphthylen-5-yl)propan-2-yl)-2-methylbenzamide (GNZL-P2)**. White solid, 20% yield. $^1$H NMR (600 MHz, DMSO-*d*6) δ 8.64 (s, 1H), 8.25 (d, J = 8.5 Hz, 1H), 7.43 (dd, J = 33.8, 7.6 Hz, 2H), 7.25 (dd, J = 21.8, 7.0 Hz, 2H), 6.77 (d, J = 8.0 Hz, 1H), 6.46 (s, 2H), 4.94 (s, 2H), 3.31 (s, 3H), 1.87 (s, 3H), 1.80 (s, 7H). $^{13}$C NMR (151 MHz, DMSO-*d*6) δ = 168.97, 146.63, 146.44, 144.66, 140.10, 139.42, 139.10, 130.94, 129.06, 127.05, 124.97, 122.13, 121.59, 119.03, 119.00, 114.69, 112.96, 55.69, 40.38, 40.24, 40.11, 39.97, 39.83, 39.69, 39.55, 30.46, 29.54, 29.45, 18.50. LC-MS (ESI, m/z): $C_{23}H_{24}N_2O$, [M+H]+ = 345.20 .

**5-amino-N-(1-(1,2-dihydroacenaphthylen-5-yl)cyclopropyl)-2-methylbenzamide (GNZL-P3).** White solid, 5% yield. $^1$H NMR (600 MHz, DMSO-*d*6) δ 8.92 (d, *J* = 2.6 Hz, 1H), 8.24 (d, *J* = 8.4 Hz, 1H), 7.70 (dd, *J* = 7.0, 2.2 Hz, 1H), 7.51 – 7.45 (m, 1H), 7.30 (d, *J* = 6.8 Hz, 1H), 7.22 (d, *J* = 7.1 Hz, 1H), 6.75 (d, *J* = 8.1 Hz, 1H), 6.43 (d, *J* = 8.1 Hz, 1H), 6.26 (d, *J* = 2.6 Hz, 1H), 3.32 (s, 4H), 3.16 (s, 2H), 1.88 (s, 3H), 1.24 (d, *J* = 7.0 Hz, 2H), 1.08 (d, *J* = 5.8 Hz, 2H). $^{13}$C NMR (151 MHz, DMSO-*d*6) δ 170.40, 146.46, 146.33, 145.48, 139.52, 138.42, 134.18, 131.01, 130.79, 130.00, 127.95, 121.49, 121.02, 119.40, 118.82, 114.97, 112.80, 33.72, 30.52, 29.88, 18.43, 14.30. LC-MS (ESI, m/z): $C_{23}H_{22}N_2O$, $[M+H]^+$ = 343.09.

**N-(1-(1,2-dihydroacenaphthylen-5-yl)cyclopropyl)-2-methyl-5-(4-methylpiperazin-1-yl)benzamide (GNZL-P4)**. White solid, 73% yield. $^1$H NMR (600 MHz, DMSO-*d*6) δ 8.99 (s, 1H), 8.27 (d, *J* = 8.4 Hz, 1H), 7.68 (d, *J* = 7.1 Hz, 1H), 7.48 (t, *J* = 7.6 Hz, 1H), 7.31 (d, *J* = 6.9 Hz, 1H), 7.23 (d, *J* = 7.1 Hz, 1H), 6.95 (d, *J* = 8.4 Hz, 1H), 6.83 (dd, *J* = 8.4, 2.6 Hz, 1H), 6.59 (d, *J* = 2.7 Hz, 1H), 3.35 (d, *J* = 7.2 Hz, 4H), 3.01 (t, *J* = 5.0 Hz, 3H), 2.45 (s, 3H), 2.23 (s, 3H), 1.95 (s, 3H), 1.32 – 1.20 (m, 4H), 1.10 (q, *J* = 4.9 Hz, 2H). $^{13}$C NMR (151 MHz, DMSO) δ 169.97, 148.95, 146.41, 145.51, 139.59, 138.13, 134.11, 131.25, 130.76, 129.84, 127.83, 125.50, 120.99, 119.42, 118.83, 116.81, 114.63, 54.87, 48.58, 45.98, 34.00, 30.54, 29.88, 18.50, 14.15. LC-MS (ESI, m/z): $C_{28}H_{31}N_3O$, [ $M+H]^+$ = 425.93.

**N-(1-(1,2-dihydroacenaphthylen-5-yl)cyclopropyl)-2-methyl-5-(piperazin-1-yl)benzamide (GNZL-P5).** Light yellow solid, 65% yield. $^1$H NMR (600 MHz, DMSO-*d*6) δ 9.55 (s, 2H), 9.09 (s, 1H), 8.27 (d, *J* = 8.3 Hz, 1H), 7.69 (d, *J* = 7.0 Hz, 1H), 7.50 (dd, *J* = 8.4, 6.8 Hz, 1H), 7.31 (d, *J* = 6.8 Hz,

1H), 7.22 (dd, *J* = 7.1, 1.6 Hz, 1H), 7.01 (d, *J* = 8.4 Hz, 1H), 6.91 (dd, *J* = 8.4, 2.6 Hz, 1H), 6.68 (d, *J* = 2.8 Hz, 1H), 3.34 (d, *J* = 7.2 Hz, 2H), 3.29 (t, *J* = 6.0 Hz, 6H), 3.15 (p, *J* = 4.5 Hz,4), 1.96 (s, 3H), 1.30 (q, *J* = 4.8 Hz, 2H), 1.11 (q, *J* = 4.8 Hz, 2H). $^{13}$C NMR (151 MHz, DMSO-$d_6$) δ 169.74, 147.77, 146.40, 145.55, 139.56, 138.37, 134.06, 131.46, 130.76, 129.91, 127.92, 127.11, 120.98, 119.46, 118.84, 117.66, 115.43, 66.82, 46.25, 42.76, 33.98, 30.53, 29.88, 18.56, 14.21. LC-MS (ESI, m/z): $C_{27}H_{29}N_3O$, $[M+H]^+$ = 412.2325.

**N-(1-(1,2-dihydroacenaphthylen-5-yl)cyclopropyl)-2-methyl-5-(4-methyl-1,4-diazepan-1-yl)benzamide (GNZL-P13).** Light yellow solid, 40% yield. $^1$H NMR (600 MHz, DMSO-$d_6$) δ 8.95 (s, 1H), 8.24 (d, *J* = 8.3 Hz, 1H), 7.67 (d, *J* = 7.0 Hz, 1H), 7.49 (dd, *J* = 8.4, 6.8 Hz, 1H), 7.31 (d, *J* = 6.9 Hz, 1H), 7.22 (d, *J* = 7.1 Hz, 1H), 6.94 (d, *J* = 8.4 Hz, 1H), 6.65 (dd, *J* = 8.5, 2.7 Hz, 1H), 6.39 (d, *J* = 2.8 Hz, 1H), 3.38 – 3.26 (m, 8H), 3.23 (s, 4H), 2.78 (s, 3H), 2.11 – 2.01 (m, 2H), 1.94 (s, 3H), 1.32 – 1.25 (m, 2H), 1.10 (q, *J* = 4.8 Hz, 2H). LC-MS (ESI, m/z): $C_{29}H_{33}N_3O$, [M+H]+ =440.23.

**N-(1-(1,2-dihydroacenaphthylen-5-yl)cyclopropyl)-5-(3-(4-hydroxy-4-methylpiperidin-1-yl)azetidin-1-yl)-2-methylbenzamide (GNZL-P17).** Brown solid, 30% yield, $^1$H NMR (600 MHz, DMSO-d6) δ 8.99 (s, 1H), 8.29 (s, 1H), 8.25 (d, J = 8.4 Hz, 1H), 7.70 (d, J = 7.0 Hz, 1H), 7.50 (t, J = 7.6 Hz, 1H), 7.31 (d, J = 6.9 Hz, 1H), 7.23 (d, J = 7.1 Hz, 1H), 6.98 (d, J = 8.2 Hz, 1H), 6.41 (d, J = 7.8 Hz, 1H), 6.20 (d, J = 2.5 Hz, 1H), 3.57-3.64 (m, 6H), 3.31 (s, 4H), 3.13 (dd, J = 7.4, 4.1 Hz, 7H), 1.96 (s, 3H), 1.60-1.69 (m, 4H).13C NMR (151 MHz, DMSO-d6) δ 12.94, 14.28, 17.18, 18.53, 18.56, 29.88, 30.53, 33.91, 39.54, 39.68, 39.82, 39.95, 40.09, 40.23, 40.37, 40.49, 42.29, 54.04, 110.87, 113.13, 118.86, 119.46, 120.96, 127.92, 130.01, 130.77, 131.21, 134.05, 138.37, 139.55, 145.57, 146.39, 169.80.LC-MS (ESI, m/z): $C_{32}H_{37}N_3O_2$, $[M+H]^+$ = 496.30.

**N-(1-(1,2-dihydroacenaphthylen-5-yl)cyclopropyl)-5-(4-(dimethylamino)piperidin-1-yl)-2-methylbenzamide (GNZL-P20).** Light yellow solid, 37% yield. 1H NMR (600 MHz, DMSO-d6) δ 9.01 (s, 1H), 8.27 (d, J = 8.4 Hz, 1H), 7.69 (d, J = 7.1 Hz, 1H), 7.49 (dd, J = 8.4, 6.8 Hz, 1H), 7.31 (d, J = 6.8 Hz, 1H), 7.23 (d, J = 7.0 Hz, 1H), 6.97 (d, J = 8.4 Hz, 1H), 6.86 (dd, J = 8.4, 2.7 Hz, 1H), 6.61 (d, J = 2.7 Hz, 1H), 3.71 – 3.63 (m, 2H), 3.35 (d, J = 7.5 Hz, 2H), 3.30 (dd, J = 9.2, 3.7 Hz, 2H), 3.08 (t, J = 11.3 Hz, 1H), 2.65 (s, 6H), 2.57 (td, J = 12.4, 2.3 Hz, 2H), 1.98 (s, 2H), 1.97 (s, 3H), 1.59 (tt, J = 12.1, 6.1 Hz, 2H), 1.29 (q, J = 4.8 Hz, 2H), 1.11 (q, J = 4.8 Hz, 2H).LC-MS(ESI, m/z): $C_{30}H_{35}N_3O$, [M+H]+ =453.93.

**N-(1-(1,2-dihydroacenaphthylen-5-yl)cyclopropyl)-2-ethyl-5-(4-methylpiperazin-1-yl)benzamide (GNZL-P25).** Light yellow solid, 23% yield. $^1$H NMR (600 MHz, DMSO-$d_6$) δ 9.14 (s, 1H), 8.39 (d, J = 7.9 Hz, 1H), 7.78 (d, J = 6.5 Hz, 1H), 7.59 (t, J = 6.9 Hz, 1H), 7.43 (d, J = 6.0 Hz, 1H), 7.34 (d, J = 6.3 Hz, 1H), 7.09 (d, J = 8.0 Hz, 1H), 6.96 (d, J = 6.9 Hz, 1H), 6.65 (s, 1H), 3.12 (s, 4H), 2.61 (s, 4H), 2.49 – 2.43 (m, 2H), 2.33 (s, 3H), 1.40 (t, J = 30.8 Hz, 6H), 1.22 (s, 2H), 0.91 (t, J = 6.9 Hz, 3H). $^{13}$C NMR (151 MHz, DMSO-$d_6$) δ 172.67, 146.42, 145.53, 145.45, 137.90, 134.24, 133.92, 130.63, 130.07, 129.87, 129.85, 127.96, 127.83, 120.93, 120.74, 119.41, 119.39, 118.83, 118.81, 115.07, 60.68, 34.04, 33.47, 32.53, 30.54, 30.51, 29.87, 29.86, 28.92, 25.17, 16.16, 14.22, 14.06.LC-MS (ESI, m/z): $C_{29}H_{33}N_3O$, $[M+H]^+$ = 440.37.

**(S)-N-(1-(1,2-dihydroacenaphthylen-5-yl)cyclopropyl)-5-(4-ethyl-3-methylpiperazin-1-yl)-2-methylbenzamide (GNZL-P28).** White solid, 10% yield. $^1$H NMR (600 MHz, DMSO-$d_6$) δ 9.00 (s, 1H), 8.27 (d, *J* = 8.4 Hz, 1H), 7.67 (d, *J* = 7.1 Hz, 1H), 7.49 (dd, *J* = 8.4, 6.8 Hz, 1H), 7.31 (d, *J* = 6.9 Hz, 1H), 7.23 (d, *J* = 7.1 Hz, 1H), 6.95 (d, *J* = 8.4 Hz, 1H), 6.82 (dd, *J* = 8.4, 2.7 Hz, 1H), 6.56 (d, *J* = 2.7 Hz, 1H), 5.75 (s, 1H), 2.61 (t, *J* = 1.9 Hz, 2H), 2.39 (t, *J* = 1.9 Hz, 2H), 1.95 (s, 3H), 1.30 – 1.27 (m, 3H), 1.26 (s, 3H), 1.24 – 1.22 (m, 4H), 1.12 – 1.09 (m, 2H), 1.01 (d, *J* = 6.0 Hz, 3H), 0.96 (t, *J* = 7.1 Hz, 3H).LC-MS (ESI, m/z): $C_{30}H_{35}N_3O$, $[M+H]^+$ = 454.28.

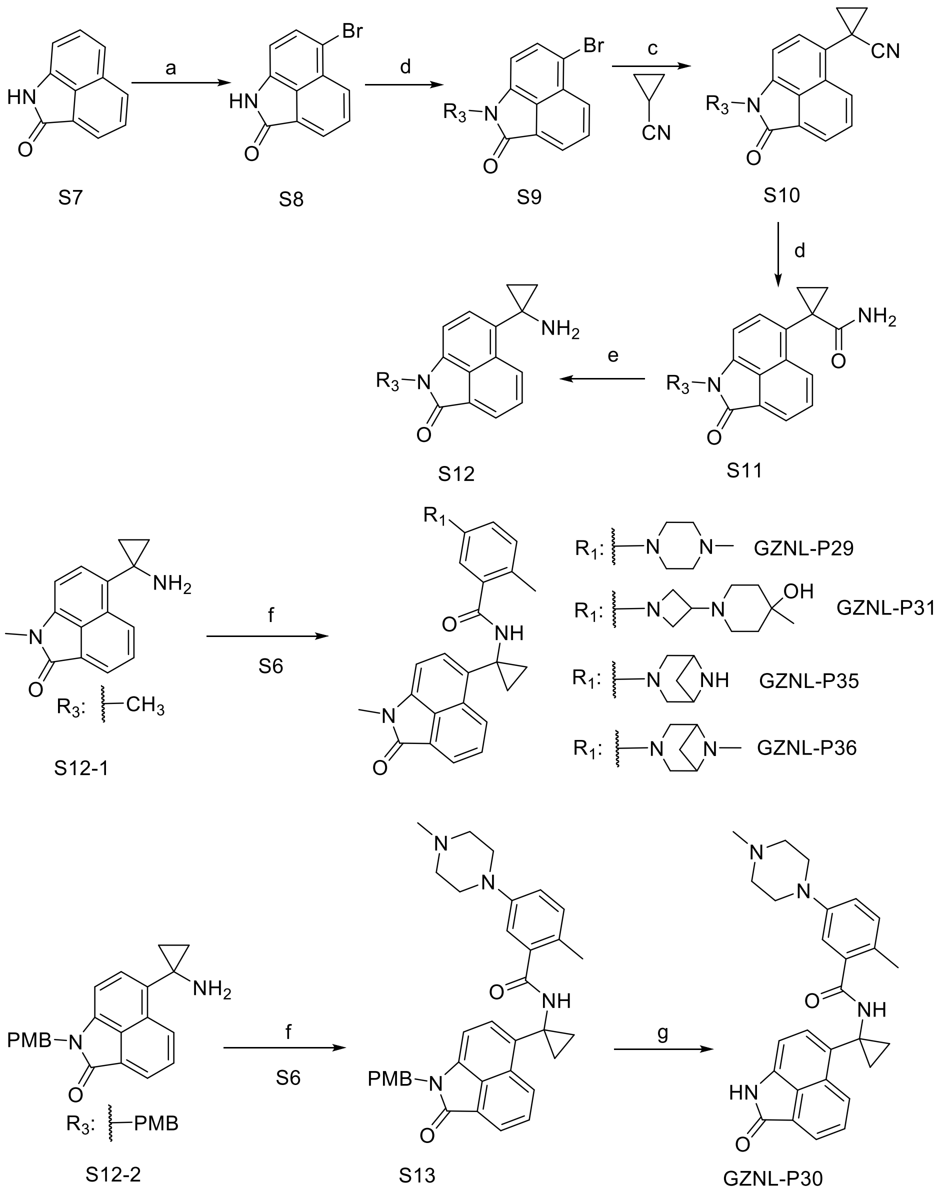


**Scheme S4. General procedure for the synthesis of compounds GZNL-P29-36.** Reagents and conditions: **a)** NBS, MeCN, 0 °C, 3 h. **b)** NaH, DMF, 0°C-rt, 1 h. **c)** LiHMDS, $Pd_2(dba)_3$, NixantPhos, THF/CPME, r.t.-60°C. **d)** $H_2O_2$, $K_2CO_3$, DMSO, 0°C-r.t., 6 h. **e)** NaClO, NaOH, *t*-BuOH, 0°C-r.t., 3 h. **f)** HATU, DIPEA, DMF, 50°C, 1 h. **g)** TFA, 60°C, overnight, 90%.

The compounds with the scaffold of benzoindolone were synthesized following the method in Scheme S4. S6 was selected as the raw material, which was converted to S7 through bromination reaction with the NBS as the brominated reagents. Substitution at $R_3$ with 4-methoxybenzyl (PMB) and methyl groups of S7 can generate the intermediate S8. This agent can be converted to S9 by coupling reaction with cyclopropanecarbonitrile. The hydrolysis reaction of the cyanide group from S9 produced compound S10, which was turned into the most important key intermediate S11 by Hofmann degradation in the presence of sodium hypochlorite/sodium hydroxide. Finally, condensation between S11 and the intermediate compound S5 generated the final products GZNL-P29, 31, 35, 36. In addition, synthesis of GZNL-P30 requires further deprotection of the PMB group after condensation.

**2-methyl-N-(1-(1-methyl-2-oxo-1,2-dihydrobenzo[cd]indol-6-yl)cyclopropyl)-5-(4-methylpiperazin-1-yl)benzamide (GNZL-P29).** Yellow solid, 81% yield. $^{1}$H NMR (600 MHz, DMSO-$d_6$) δ 9.06 (s, 1H), 8.81 (d, $J$ = 8.2 Hz, 1H), 8.04 (d, $J$ = 6.9 Hz, 1H), 7.83 (dd, $J$ = 8.3, 7.0 Hz, 1H), 7.73 (d, $J$ = 7.3 Hz, 1H), 7.08 (d, $J$ = 7.3 Hz, 1H), 6.96 (d, $J$ = 8.4 Hz, 1H), 6.84 (dd, $J$ = 8.4, 2.7 Hz, 1H), 6.59 (d, $J$ = 2.7 Hz, 1H), 3.36 (s, 3H), 3.03 (t, $J$ = 5.0 Hz, 4H), 2.27 (s, 3H), 1.93 (s, 3H), 1.36 – 1.28 (m, 3H), 1.24 (d, $J$ = 9.3 Hz, 3H), 1.20 – 1.17 (m, 2H). $^{13}$C NMR (151 MHz, DMSO-$d_6$) δ 170.00, 167.74, 139.35, 133.02, 131.31, 130.40, 130.10, 128.90, 128.63, 126.92, 125.22, 124.09, 116.90, 114.47, 105.34, 33.72, 26.61, 18.43, 14.16. LC-MS (ESI, m/z): $C_{28}H_{30}N_4O_2$, $[M+H]^+$ = 454.37.

**2-methyl-5-(4-methylpiperazin-1-yl)-N-(1-(2-oxo-1,2-dihydrobenzo[cd]indol-6-yl)cyclopropyl)benzamide (GNZL-P30).** Yellow solid, 55% yield. $^{1}$H NMR (600 MHz, DMSO-$d_6$) δ 10.73 (s, 1H), 9.05 (s, 1H), 8.80 (d, $J$ = 8.2 Hz, 1H), 8.00 (d, $J$ = 6.8 Hz, 1H), 7.83 (t, $J$ = 7.4 Hz, 1H), 7.68 (d, $J$ = 7.1 Hz, 1H), 6.99 (d, $J$ = 8.0 Hz, 1H), 6.91 (d, $J$ = 7.2 Hz, 1H), 6.87 (d, $J$ = 7.9 Hz, 1H), 6.62 (s, 1H), 3.12 (s, 3H), 2.83 (s, 3H), 1.95 (s, 3H), 1.34 (d, $J$ = 9.3 Hz, 1H), 1.30 (s, 2H), 1.23 (s, 4H), 1.18 (s, 2H). $^{13}$C NMR (151 MHz, DMSO-$d_6$) δ 169.88, 169.48, 138.17, 137.86, 132.30, 131.38, 130.56, 130.01, 129.06, 128.86, 127.71, 126.54, 126.05, 123.79, 114.74, 105.93, 47.51, 33.70, 29.45, 18.45, 14.19. LC-MS (ESI, m/z): $C_{27}H_{28}N_4O_2$, $[M+H]^+$ = 441.22.

**5-(3-(4-hydroxy-4-methylpiperidin-1-yl)azetidin-1-yl)-2-methyl-N-(1-(1-methyl-2-oxo-1,2-dihydrobenzo[cd]indol-6-yl)cyclopropyl)benzamide (GNZL-P31).** Yellow solid, 61% yield. $^{1}$H NMR (600 MHz, DMSO-$d_6$) δ 9.04 (s, 1H), 8.79 (d, $J$ = 8.1 Hz, 1H), 8.16 (s, 1H), 8.03 (d, $J$ = 6.7 Hz, 1H), 7.82 (t, $J$ = 7.4 Hz, 1H), 7.72 (d, $J$ = 7.1 Hz, 1H), 7.07 (d, $J$ = 7.1 Hz, 1H), 6.91 (d, $J$ = 8.0 Hz, 1H), 6.33 (d, $J$ = 7.1 Hz, 1H), 6.12 (s, 1H), 3.83-3.80 (m, 2H), 3.41-3.39 (m, 2H), 3.35 (s, 3H), 3.20-3.17 (m, 1H), 2.27 (d, $J$ = 47.1 Hz, 4H), 1.90 (s, 3H), 1.44 (t, $J$ = 16.7 Hz, 4H), 1.29 (s, 2H), 1.18 (s, 2H), 1.08 (s, 3H). $^{13}$C NMR (151 MHz, DMSO-$d_6$) δ 170.06, 167.74, 150.13, 139.34, 137.97, 133.02, 131.13, 130.43, 130.10, 128.91, 125.18, 124.09, 123.21, 112.74, 110.36, 105.35, 66.05, 56.87, 54.97, 38.44, 33.66, 26.60, 18.46, 14.19. LC-MS (ESI, m/z): $C_{32}H_{36}N_4O_3$, $[M+H]^+$ = 525.25.

**5-(3,6-diazabicyclo[3.1.1]heptan-3-yl)-2-methyl-N-(1-(1-methyl-2-oxo-1,2-dihydrobenzo[cd]indol-6-yl)cyclopropyl)benzamide (GNZL-P35).** Yellow solid, 82% yield. $^{1}$H NMR (600 MHz, DMSO-$d_6$) δ 9.05 (s, 1H), 8.82 (d, $J$ = 8.2 Hz, 1H), 8.04 (d, $J$ = 6.9 Hz, 1H), 7.83 (dd, $J$ = 8.2, 7.0 Hz, 1H), 7.73 (d, $J$ = 7.3 Hz, 1H), 7.09 (d, $J$ = 7.3 Hz, 1H), 6.98 (d, $J$ = 8.5 Hz, 1H), 6.62 (dd, $J$ = 8.4, 2.7 Hz, 1H), 6.39 (d, $J$ = 2.7 Hz, 1H), 3.99 (d, $J$ = 5.4 Hz, 2H), 3.49 (d, $J$ = 11.0 Hz, 2H), 3.36 (s, 3H), 2.65 – 2.61 (m, 1H), 1.95 (s, 3H), 1.57 (d, $J$ = 9.1 Hz, 1H), 1.32 – 1.30 (m, 2H), 1.23 (s, 2H), 1.21 – 1.19 (m, 2H). $^{13}$C NMR (151 MHz, DMSO-$d_6$) δ 170.40, 167.74, 147.10, 139.33, 138.09, 133.09, 131.32, 130.31, 130.12, 128.89, 128.66, 126.91, 121.75, 111.16, 109.03, 105.32, 55.58, 51.10, 33.76, 18.36, 14.17. LC-MS (ESI, m/z): $C_{28}H_{28}N_4O_2$, $[M+H]^+$ = 453.39.

**2-methyl-N-(1-(1-methyl-2-oxo-1,2-dihydrobenzo[cd]indol-6-yl)cyclopropyl)-5-(6-methyl-3,6-diazabicyclo[3.1.1]heptan-3-yl)benzamide (GNZL-P36).** Yellow solid, 77% yield. $^{1}$H NMR (600 MHz, DMSO-$d_6$) δ 9.09 (s, 1H), 8.83 (d, $J$ = 8.2 Hz, 1H), 8.04 (d, $J$ = 6.8 Hz, 1H), 7.84 (t, $J$ = 7.5 Hz, 1H), 7.74 (d, $J$ = 7.2 Hz, 1H), 7.09 (d, $J$ = 7.2 Hz, 1H), 7.00 (d, $J$ = 8.2 Hz, 1H), 6.64 (d, $J$ = 7.4 Hz, 1H), 6.42 (s, 1H), 3.89 (s, 2H), 3.36 (s, 3H), 2.50 (s, 3H), 2.10 (s, 2H), 1.95 (s, 3H), 1.62 (s, 1H), 1.34 (d, $J$ = 9.3 Hz, 1H), 1.31 (d, $J$ = 9.2 Hz, 2H), 1.26 (s, 2H), 1.20 (s, 2H). $^{13}$C NMR (151 MHz, DMSO-$d_6$) δ 170.26, 167.74,

145.71, 139.34, 138.21, 133.05, 131.42, 130.37, 130.12, 128.93, 128.65, 126.90, 125.21, 124.11, 105.35, 33.75, 30.85, 29.45, 26.61, 22.57, 18.33, 14.19. LC-MS (ESI, m/z): $C_{29}H_{30}N_4O_2$ $[M+H]^+ = 467.32$.

### LogD and solubility

To test the partition coefficient between the aqueous and organic phase (LogD) of GZNL-P36, 15 µL of stock solutions (10 mM) of each sample were placed into their proper 96-well rack, 500 µL of PBS saturated 1-octanol was added into each vial of the cap-less LogD plate followed by the addition of 500 µL of 1-octanol saturated PBS (*p*H 7.4). One stir stick was added to each vial and a molded PTFE/Silicone plug was used to seal each vial. Then the LogD plate was transferred to the Eppendorf Thermomixer Comfort plate shaker and shaken at 25 °C with 1,100 RPM for 1 hour. After completion of 1 hour, plugs were removed and the stir sticks were removed using a big magnet. The samples were then centrifuged at 25°C at 20,000 *g* for 20 min to separate the phases, and pipette and syringe were used to remove the upper (1-octanol) and lower (buffer) phases to the empty tubes, respectively. Aliquots of 5 µL were taken from the upper phases followed by an addition of 495 µL of a mixture of $H_2O$ and acetonitrile (1:1 in v/v). Vortex for 1 minute, and then aliquots of 50 µL were taken from the diluent followed by an addition of 450 µL of a mixture of $H_2O$ and acetonitrile (1:1 in v/v). Then, aliquots of 50 µL were taken from lower phases followed by an addition of 450 µL of a mixture of $H_2O$ and acetonitrile (1:1 in v/v). 200 µL of diluent was transferred to a new 96-well plate for LC-MS/MS analysis.

To test the solubility of GZNL-P36, a proper amount of compounds was weighed out and placed into a 96-well rack. Based on the amount, the proper volume of PBS *p*H 7.4 was added to each vial of the solubility sample plate which was transferred to the Eppendorf Thermomixer Comfort plate shaker and shaken at 25°C and 1,100 rpm for 24 h. After completion of 24 h, the samples were transferred from the Solubility Sample plate into a filter plate for removing the un-soluble ingredient using the vacuum manifold. The filtrate was diluted by 1000-fold ((An aliquot of 5 µL was taken from the filtrate followed by the addition of 5 µL DMSO and 490 µL of a mixture of $H_2O$ and acetonitrile (1:1 in v/v), vortex well and then aliquot of 50 µL was taken from the diluent followed by addition of 450 µL of a mixture of $H_2O$ and acetonitrile (1:1 in v/v)). 200 µL of diluent was transferred to a new 96-well plate for LC-MS/MS analysis.

### Analysis of compound metabolites

The *in vitro* metabolites analysis of GZNL-P4 was evaluated using rat liver cells. The compound was diluted to 20 µM using William's E medium as a test solution. Rat liver cell suspension was prepared at the density of $1 \times 10^6$ cells/mL. The test solution and cell suspension were incubated in a constant temperature oscillation incubator at 37 °C with 5% $CO_2$ for 5 min. For the time point 0 min, 600 µL of acetonitrile and 100 µL of cell suspension were mixed by vortex oscillation for 5 min, then followed by 100 µL of the test solution and another vortex oscillation for 5 min. For the time point 20 min, 100 µL of test solution was added into 100 µL of cell suspension to start the reaction. After incubating in a constant temperature oscillation incubator at 37 °C with 5% $CO_2$ for 20 min, 600 µL of acetonitrile was added into the sample and oscillated by vortex for 5 min. All samples were centrifuged at 18,000 × *g* for 10 min, and the supernatant was transferred to fresh tubes for use. 700 μL of supernatant was dried under nitrogen airflow at 37 °C. The dried sample was re-dissolved by 200 μL of 75% acetonitrile (v/v) for LC-UV-HRMS analysis (Vanquish Flex, Q Exactive HF-X, Thermo Scientific).

**Metabolic stability in liver microsome**

The metabolism in human liver microsomes was used to evaluate the *in vitro* metabolic stability of GZNL-P4, GZNL-P17, GZNL-P29, GZNL-P31, GZNL-P35, and GZNL-P36. 1.5 μM of compounds were incubated with rat or human liver microsomes (0.75 mg/mL) in a volume of 0.5mL PBS, respectively. Assay plates dispensed 30 μL of 1.5 μM compound solution containing 0.75 mg/mL microsomes were designated for different time points (0, 5, 15, 30, and 45 min). NADPH (final concentration at 2 mM) was used to start the reaction at 37 °C and acetonitrile solution containing IS (100 nM alprazolam, 200 nM labetalol, 200 nM caffeine and 2 μM ketoprofen) was used to terminate the reaction at different time points. The plates were centrifuged at 6000 rpm for 15 min after shaking for 10 min at 600 rpm. Precipitated protein was removed and 80 μL supernatant was transferred to 96-well plate containing 140 μL pure water for LC/MS (Shimadzu Nexera LC-40 & SCIEX TQ-6500+) analysis. Intrinsic clearance by substrate depletion was calculated taking into account mg of microsomes per liver weight and grams of liver per Kg body weight.

**Metabolic stability in hepatocyte**

The cryopreserved hepatocytes were thawed in a 37°C water bath and gently shaking the vials for 2 min. The hepatocytes were transferred into 50 mL conical tube containing thawing medium. The thawing medium was removed after centrifuged at 100 × *g* for 10 minutes and the hepatocytes were resuspended in enough incubation medium to yield $1.5 \times 10^6$ cells/mL. 1 μM compound was incubated with pre-warmed hepatocytes in incubation medium at $0.5 \times 10^6$ live cells/mL. The 25 μL sample taken at time points of 0, 15, 30, 60, 90 and 120 minutes, were mixed with 6 volumes (150 μL) of acetonitrile containing internal standards (IS: 100 nM alprazolam, 200 nM labetalol, 200 nM caffeine and 2 μM ketoprofen) to terminate the reaction. After centrifugation for 20 min at 3,220 × *g*, aliquot of 100 μL of the supernatant was mixed with 100 μL of ultra-pure water for LC-MS/MS analysis. Boiled cells were used as control. The *in vitro* half-life (*in vitro* $T_{1/2}$) was determined from the slope value:

*in vitro* $T_{1/2} = 0.693/k$,

where k = rate constant (-slope value).

The intrinsic clearance (CLint, in μL/min/$10^6$cells) was calculated using the following equation:

CLint = kV/N,

where V = incubation volume (0.2 mL);

N = number of hepatocytes per well ($1 \times 10^5$ cells)

**Inhibition of hERG**

To evaluate the inhibitory effects of GZNL-P36 on the hERG, the current on human embryonic kidney cells stably expressing hERG channels (hERG-HEK293 cells, Creacell, A-0320) was monitored using a patch clamp system that the core components were bought from Sutter Instrument (USA). The cells were clamped using the patch clamp system to perform a whole-cell voltage clamp mode, the hERG currents were elicited with corresponding voltages. The cells were treated with serially diluted cisapride (sigma, positive control) and GZNL-P36. Tail currents of hERG channels were recorded to obtain the peak tail current at each concentration using Patchcontrol HT and Patchmaster. The current recorded at non-compound containing solution was used as control for each cell. The current was recorded twice independently for each cell, and at least 2 cells were recorded or each concentration.

**Inhibition of CYP isoforms**

In this study, test compounds were added into the liver microsomes solution pre-warmed at 37 °C for 5 min. Subsequently，20 μL of pre-warmed (37 °C) 10 mM NADPH solution was added into 180 μL of liver microsomes solution treated by compounds to initiate the enzyme reaction. The assay plate was incubated at 37 °C until to the designated time when 300 μL of termination solution (35 ng/mL ketoprofen, 7.5 ng/mL carbamazepine, 5 ng/mL diphenhydramine, 10 ng/mL tolbutamide) was added to quench the reaction. Then the samples were centrifuged at 3,220 × g for 40 min. The supernatant was transferred to analysis plate containing a proper volume of ultra-pure water for LC-MS/ MS analysis. The inhibition IC50 was determined for the CYP isoforms 1A2, 2C9, 2C19, 2D6 and 3A4.

**Plasma protein binding assay**

Firstly, a basic solution was prepared by dissolving 14.2 g/L $Na_2HPO_4$ and 8.77 g/L NaCl in deionized water and the solution could be stored at 4°C for up to 7 days. And an acidic solution was prepared by dissolving 12.0 g/L $NaH_2PO_4$ and 8.77 g/L NaCl in deionized water and the solution could be stored at 4°C for up to 7 days. Then, the working buffer was prepared by titrating the basic solution with the acidic solution to *p*H 7.4 and store at 4°C for up to 7 days. *p*H was checked on the day of experiment and was adjusted if outside specification of 7.4 ± 0.1. The plasma stored at −80 °C was thawed and centrifuged at 3,220 ×*g* for 10 min to remove clots and the supernatant was collected into a fresh tube. Compounds were added into the plasma to achieve a final concentration of 1 μM (0.5% DMSO). The dialysis membranes were soaked in ultra-pure water for 60 min to separate strips, then in 20% ethanol for 20 minutes, finally in dialysis buffer for 20 min. 50 μL of the spiked plasma samples were transferred to separate wells of a 96-well plate and the plate was incubated at 37°C with 5% $CO_2$ for 6 hours. At the end of incubation, samples from both buffer and plasma chambers were transferred to wells of a 96-well plate. A room temperature quench solution (acetonitrile containing 200 nM labetalol, 100 nM tolbutamide and 100 nM ketoprofen) was added to precipitate protein. Samples in plate were vortexed and centrifuged at 3,220 × *g* for 30 min at

4°C. And 100 μL of supernatant was transferred to a new 96-well plate with 100 μL of water for LC-MS/MS analysis.

**In vivo PK experiments**

In this study, mouse and dog pharmacokinetic studies were done at Medicilon (Shanghai, PRC); male ICR mice (n = 3) and male beagle dogs (n = 3) were treated with intravenous or oral administration. The vehicle used for the ICR mice consisted of 5% DMSO, 10% solution, and 85% saline. Blood samples were collected at 0.25h, 0.5h, 1h, 2h, 4h, 6h, 8h, 10h, 12h, 24h after administration and analyzed using liquid chromatography tandem mass spectrometry (LC-MS/MS). The pharmacokinetic parameters $AUC_{0\text{-}t}$、 $AUC0_{\text{-}\infty}$、 $MRT0_{\text{-}\infty}$、 $C_{max}$、 $T_{max}$、 $T_{1/2}$ were calculated using Phoenix WinNonlin 7.0.

**Plasmid Construction, Protein Expression and Purification**

The papain-like protease domain sequence was obtained from the SARS-CoV-2 complete genome (NCBI gene databank, severe acute respiratory syndrome coronavirus 2, Gene ID: 43730578). The $PL^{pro}$ Ubl domain protein sequence (amino acids, 746-1060) of Nsp3 protein from SARS-CoV-2 (Nsp3, YP_009742610.1) was codon optimized, synthesized, and cloned into pET28a with BamHI and XhoI (Tsingke). The TEV protease recognition sequence was inserted between the N-terminal $His_6$-tag and $PL^{pro}$ sequence. The $PL^{pro}$ mutant C111S ($PL^{proC111S}$) was generated by PCR. BL21(DE3) *Escherichia coli* competent cells were used for protein expression. The competent cells transformed with $PL^{pro}$ expression plasmid were inoculated in LB medium and grown to an OD600 of 0.6-0.8, 0.5 mM of isopropyl-D-thiogalactopyranoside (IPTG), and 1mM zinc chloride (ZnCl2) were used to induce protein production at 16°C for 18 hours. Cell pellets were collected by centrifugation at 4000 rpm for 30 minutes and then stored at −80 °C for use. The cell pellets were thawed and resuspended with buffer A [50 mM Tris-HCl, 150 mM NaCl, 10 mM imidazole, and 1 mM TCEP (*p*H 7.4)] supplemented with lysozyme, deoxyribonuclease, and cOmpleteTM protease inhibitor cocktail (Roche). After lysed by sonication, the cell lysates were separated by centrifugation at 18,000 × *g* for 35 minutes twice at 4 °C. The supernatant was further filtered using a 0.22 μm pore size membrane. His-tagged proteins were loaded onto a HisTrap HP column (5 ml; GE) equilibrated with buffer A for 10 column volumes. The unspecific binding proteins were washed with buffer A for 16 column volumes and the target protein was eluted with buffer B [50 mM Tris-HCl, 150 mM NaCl, 250 mM imidazole, and 1 mM TCEP (*p*H 7.4)] for 20 column volumes by linear gradient. The His-tag was removed by TEV protease during dialysis. Then, the protein cleaved His-tag was concentrated to 2 ml and loaded onto Superdex 75 column (120 ml; GE) equilibrated with buffer C [20 mM Tris-HCl, 100 mM NaCl, and 1 mM TCEP *(p*H 7.4)] for 1 column volumes.

The recombinant expression plasmids of other $PL^{pro}$s were constructed following the same procedure of SARS-CoV-2 $PL^{pro}$. The protein expression and purification of these $PL^{pro}$s are also following the similar protocol of SARS-CoV-2 $PL^{pro}$ with minus changes. The $PL^{pro}$s tested in this study are from four different subfamily coronaviruses, including alpha-coronavirus (TEGV-$PL^{pro}$, Mink-$PL^{pro}$, Feline-$PL^{pro}$, HCoV-NL63-$PL^{pro}$, HCoV-229E-$PL^{pro}$), beta-coronavirus (HCoV-SARS-PL$^{pro}$, HCoV-SARS-2-PL$^{pro}$, Equine-PL$^{pro}$,), delat-coronavirus (pigeon-PL$^{pro}$, PDCoV-PL$^{pro}$, quail-PL$^{pro}$), and gamma- coronavirus (Avian-PL$^{pro}$).

**PL$^{pro}$ enzymatic assay**

The activity of PL$^{pro}$ was measured using the labeled peptide Z-RLRGG-AMC (GLPBIO, GA23715) as substrate in a black 384-well plate (GREINER, 784076), using wavelengths of 340 nm and 460 nm for excitation and emission, respectively. The assay buffer contains 50mM HEPES, 10mM DTT, 0.1mM EDTA, 0.005% tween 20, *p*H7.2. The gradient diluted compounds were added into PL$^{pro}$ solution diluted with assay buffer and incubated for 10 minutes. The substrate peptide diluted with assay buffer was added into PL$^{pro}$ and inhibitor mix solution and incubated for 1 hour at 37 °C. The final concentrations are 10 nM, 20 μM for PL$^{pro}$ and substrate, respectively. Fluorescence intensity was monitored with a BioTek Neo2 multimode plate reader (Agilent). The results were plotted as dose inhibition curves using nonlinear regression to determine the $IC_{50}$ values of inhibitor compounds using GraphPad Prism 9.0. The enzymatic inhibition efficacy of GZNL-P36 against PL$^{pro}$s was calculated:

Efficacy = (Top – Bottom)/Top,

Where Top and Bottom are taken from the corresponding terms in the nonlinear fitting formula,

Then, all the efficacies of PL$^{pro}$s are normalized to that of SARS-CoV-2 PL$^{pro}$.

**Crystallization, data collection and structure determination**

Purified SARS-CoV-2 PL$^{proC111S}$ was incubated with inhibitors (GZNL-P4, GZNL-P28, GZNL-P31, and GZNL-P35) at a 1:3 protein/ligand molar ratio in buffer overnight and concentrated to 8 mg/ml. Protein-ligand complex crystals were obtained after 7-10 days at 4 °C by sitting-drop vapor diffusion against 40 μl of well solution using 96-well crystallization plates. The crystallization drops contained 0.75 μL of protein complex sample mixed with 0.75 μL of reservoir solution. The crystals with well diffraction ability were obtained in conditions containing 0.1 M HEPES *p*H 7.5, 0.2 M Lithium sulfate monohydrate, and 25% (w/v) Polyethylene glycol 3350 for GZNL-P4; 0.03 M Citric acid, 0.07 M BIS-TRIS propane *p*H 7.6, 20% w/v Polyethylene glycol 3,350 for GZNL-P28; 5mM $CoCl_2$-6$H_2O$, 5mM $NiCl_2$-6$H_2O$, 5mM $CdCl_2$-$H_2O$, 5mM $MgCl_2$-6$H_2O$, 0.1 M HEPES *p*H 7.5, 12% w/v PEG3,350 for GZNL-P31; 0.1 M Bis-Tris *p*H 6.5, 0.2 M Magnesium chloride hexahydrate, and 25% (w/v) Polyethylene glycol 3350 for GZNL-P35.

X-ray diffraction data were collected at beamline BL19U1 and BL02U1 at the Shanghai Synchrotron Radiation Facility and processed with the program autoPROC1.0[43]. The structures were solved by molecular replacement using the program Phaser[44] integrated in PHENIX[45] package using the previously published PL$^{pro}$ structure (PDB code: 7CJD)[46] as the search model. The structures were refined using PHENIX with several cycles of manually interactive model rebuilding in Coot[47]. The ligand constraints were generated using Phenix.elbow[48] and the ligand models were built according to the omit map. The data collection and refinement statistics are summarized in **Extended Data Table S2**.

**Bio-layer interferometry (BLI) assay.**

BLI assay was carried out at 25 °C using Octet R8 (Sartorius) with phosphate buffer solution (PBS) containing 0.005% tween-20 (PBST). Biotinylation SARS-CoV-2 $PL^{pro}$ protein was produced using a commercially available kit (G-MM-IGT, Genemore). The $PL^{pro}$ protein (50 μg/mL) was immobilized onto Super Streptavidin SSA capture biosensors (18-5070, Sartorius) pre-equilibrated with PBST. The biosensors were then exposed to different concentrations of compound prepared with PBST for association, followed by the dissociation in PBST. Data was analyzed using the Octet® BLI Analysis software.

**Isothermal titration calorimetry (ITC) assay**

ITC assays were carried out at 25 °C with 19 injections including the first pre-injection of 0.4 μL and 18 injections of 2 μL using a MicroCal PEAQ-ITC instrument (Malvern Panalytical). Reference power was set to 5 μcal/s, and the compound solution was titrated at 150 s. The $PL^{pro}$ proteins and compounds were diluted with a buffer (*p*H 7.4) containing 20 mM Tris-HCl, 100 mM NaCl, 1 mM TECP, and 0.5% (v/v) DMSO. All samples were centrifugated at 13000 rpm for 10 minutes to remove precipitation and bubbles before titration. The $PL^{pro}$ proteins (20 μM) were injected into the sample cell and compounds (200 μM) were loaded into the syringe cell, whereas deionized water was injected into the reference cell as a heat balance control. Data were fitted into a one-site model, as well as $K_d$, N, ΔG, ΔH, and −TΔS values were calculated using MicroCal PEAQ-ITC analysis software.

**Cell culture and cell viability assay**

Vero E6 (ATCC, CRL-1586), Huh-7 (JCRB, 0403), HEK293T (ATCC CRL-3216) and HCT-8 (ATCC, CCL-244) cells were cultured in Dulbecco's Modified Eagle Medium (DMEM) supplemented with 10% fetal bovine serum (FBS), 100 IU/mL penicillin and 100 μg/mL streptomycin. The cells were cultured at 37°C in a fully humidified atmosphere containing 5% $CO_2$, and have been tested negative for mycoplasma infection.

Cell viability was evaluated using a Cell Titer-Glo 2.0 Cell Viability Assay (G9242, Promega) according to the manufacturer's instructions. In brief, $4 \times 10^3$ cells in 20 μL culture medium were seeded into opaque-walled 384-well plates and incubated for 12 hours for adherence. After being treated with compounds for 48 h, 20 μL of Cell Titer-Glo reagent was added into each well. After 2 min shaking and 10 min incubation, luminescence was measured by BioTek Synergy H1 multimode reader (Agilent).

**Virus preparation and titrations**

SARS-CoV-2 wild-type strain (WT, GenBank: MT123291), Omicron BA.5 variant (Omicron BA.5, GDPCC-303), and XBB.1 variant (XBB.1, IQTC-1596943) were propagated in Vero E6 cells and stored at −80 °C. Virus titers were determined with 10-fold serial dilutions in confluent Vero E6 cells in 96-well microtitre plates. Three days after inoculation, a cytopathic effect (CPE) was scored, and the Reed-Muench formula was used to calculate the $TCID_{50}$. All of the infection experiments were performed at BSL-3 in Guangzhou Customs Inspection and Quarantine Technology Center (IQTC).

HCoV-OC43 strain (VR-1558) and HCoV-NL63 strain (NR-470) were propagated in HCT-8 and Huh7-hACE2 cells and stored at −80 °C, respectively. Virus titers were determined with an indirect immunofluorescence assay. In brief, the cells were fixed in 4% paraformaldehyde for 15 minutes at 24 hours post-infection and permeabilized with 0.3% Triton X-100 for 15 minutes at room temperature. HCoV-OC43 and HCoV-NL63 were probed using anti-HCoV-OC43 Nucleoprotein antibody (Abcam, ab309964, 1:100) and anti-HCoV-NL63 Nucleoprotein antibody (SinoBiological, 40641-T62, 1:500) over 1 hour incubation at room temperature and followed by Alexa Fluor488-labelled secondary antibody (Jackson), respectively. All the cells were stained with 4,6-diamidino-2-phenylindole (DAPI, Sigma, USA) for nuclear visualization. Fluorescent images were acquired using an Evos M5000 Cell Imaging System (Thermo Fisher Scientific, Massachusetts, USA) at 4×magnification. Fluorescence dots were quantified and normalized to the total nuclear count using Image J. All of the infection experiments were performed at BSL-2 in Guangzhou Laboratory.

#### *In vitro* antiviral activity assay

$2 \times 10^4$ Vero E6 cells were seeded in a 96-well plate for 24 hours. The compounds with different dilution concentrations were mixed with SARS-CoV-2 (MOI=0.01), and 200 μL mixtures were inoculated onto monolayer Vero E6 cells. Seventy-two hours after inoculation, CPE was scored by Celigo Image Cytometer. The inhibition of compounds and the value of $EC_{50}$ is calculated from SARS-CoV-2 's CPE rates. Three independent experiments were performed with eight concentration gradients, each with triplicate wells, and one representative is shown.

$2 \times 10^4$ Huh-7 cells were seeded in a 96-well plate for 24 h. GZNL-P36 with different dilution concentrations were mixed with HCoV-229E (MOI=0.3), and 200 μL mixtures were inoculated onto monolayer Huh-7 cells. Forty-eight hours after inoculation, CPE was scored by Celigo Image Cytometer. The inhibition of compounds and the value of $EC_{50}$ are calculated from HCoV-229E's CPE rates. Three independent experiments were performed with eight concentration gradients, each with triplicate wells, and one representative is shown.

$1 \times 10^5$ Huh-7 (for HCoV-OC43) or Huh7-hACE2 (for HCoV-NL63) cells were seeded in a 24-well plate for 24 h, respectively. GZNL-P36 with different dilution concentrations were mixed with HCoV-OC43 (MOI=0.3) or HCoV-NL63 (MOI=0.3), and 200 μL mixtures were inoculated onto monolayer cells. Forty-eight hours after inoculation, the intracellular total RNA was collected by Trizol reagent. The viral RNA level was determined using qRT-PCR. For HCoV-OC43, the primer (F/R, 5’-3’) targets N gene: GCTCAGGAAGGTCTGCTCC/TCCTGCACTAGAGGCTCTGC. For HCoV-NL63, the primer (F/R, 5’-3’) targets N gene: AGGACCTTAAATTCAGACAACGTTCT/GATTACGTTTGCGATTACCAAGACT, probe: 5'-FAM-TAACAGTTTTAGCACCTTCCTTAGCAACCCAAACA-3'-BHQ1. The inhibition rates were calculated as the percentage of the viral RNA level relative to the control. Two independent experiments were performed with eight concentration gradients, each with triplicate wells, and one representative is shown.

**SARS-CoV-2 infection in K18-hACE2 mice**

K18-hACE2 transgenic mice aged 8 weeks were obtained from the GemPharmatech. The use of K18-hACE2 transgenic mice has received ethical approval from the Animal Ethics Committee at Guangzhou Customs Inspection and Quarantine Technology Center (IQTC2023015). Forty-eight female hACE2 transgenic mice were divided into six groups with eight mice in each group to evaluate the efficacy of GZNL-P36 in the therapeutic treatment. On the day of infection, the hACE2 mice were intranasally inoculated with either $5\times10^4$ $TCID_{50}$ XBB.1, pre-diluted in 50 μL DMEM. Treatment was delayed until 2 hours post-infection (h.p.i.). K18-hACE2 transgenic mice were orally administered a dose of 25 mg/kg S-217622, GZNL-P36 (25, 50 or 100 mg/kg) diluted in 200 μL 5% DMSO/20% hydroxypropyl-beta-cyclodextrin for the treatment group or vehicle solution only for the control group. Mice were killed at the designated time points and organ tissues were sampled for virological and histopathological analysis.

**RNA extraction and qRT-PCR**

Tissue samples were lysed and extracted with the Trizol (Invitrogen) reagent according to the manufacturer's protocols. After RNA extraction, complementary DNA was synthesized through reverse transcription using the Evo M-MLV RT Mix Kit with gDNA Clean reagent for qPCR (AG11705, Accurate Biology). Real-time PCR was carried out using 2 × SYBR Green *Pro Taq* HS Master Mix qPCR Kit (AG11701, Accurate Biology) on a CFX384 Touch Real-Time PCR Detection System (Bio-Rad). The setting procedures were as follows: 95°C for 30 seconds, 95°C for 5 seconds, 60°C for 30 seconds, a total of 40 cycles. Glyceraldehyde-3-phosphate dehydrogenase (GAPDH) was used as an internal control. The relative expression level of the target gene was calculated by the $2^{-\Delta\Delta CT}$ method. The specific primers of immune-related genes used for Real-time PCR are listed as follows:

*GAPDH* sense (5'- AATGAAGGGGTCATTGATGG -3') and antisense (5'-AAGGTGAAGGTCGGAGTCAA -3'),

*IFNB1* sense sense (5'- AGCTCCAAGAAAGGACGAACA-3') and antisense (5'-GCCCTGTAGGTGAGGTTGAT-3'),

*CXCL10* sense (5'-CCAAGTGCTGCCGTCATTTTC -3') and antisense (5'-GGCTCGCAGGGATGATTTCAA-3'),

*IFNgamma1* sense (5'-GCCACGGCACAGTCATTGA-3') and antisense (5'-TGCTGATGGCCTGATTGTCTT-3').

**Fluorescence focus assay**

Confluent monolayers of Vero E6 cells were incubated with 50 μL pulmonary tissue homogenate of mice with threefold serial dilutions for 2 h at 37 °C, 5% $CO_2$, in triplicate per condition. The inoculum was removed and cells were overlayed with a virus growth medium containing 2% carboxymethylcellulose sodium. At 24 hours post-infection, cells were fixed in 4% paraformaldehyde and permeabilized with 0.2% Triton-X-100, and virus plaques were visualized by immunostaining using an anti-nucleocapsid antibody and action of HRP on a tetramethylbenzidine-based substrate.

**Histology and immunohistochemistry**

Hematoxylin/eosin (HE) staining of lung samples followed standard HE staining procedures. The lung tissue was cut off and fixed in a 4% polymethylaldehyde solution for 24 hours. Briefly, the sections (3-4 μm-thick) were stained with hematoxylin for 3 minutes and then soaked in the acidic liquid alcohol differentiation for 30 s. After staining with eosin for 15 s and dehydrated by ethanol, the sections were finally cleared by xylene and mounted. The images of per slide were captured using a microscope (LEICA Aperio Versa 8, Germany) and analyzed by the Image J software.

The lung tissue was dewaxed in xylene and hydrated in ethanol. The endogenous peroxidase was blocked using a 3% $H_2O_2$ solution. The antigen was retrieved by boiling in a citric acid solution (*p*H=6.0), and non-specific binding sites were blocked with 3% BSA at room temperature for 30 minutes. The tissues were then incubated with anti-SARS-CoV-2 (COVID-19) Nucleocapsid primary antibodies (1:1,000 dilution) at 4°C overnight. After washing, the sections were conjugated with a horseradish peroxidase (HRP) antibody (1:200 dilution; Servicebio) at room temperature for 50 minutes. The tissues were developed using 3,3'-diaminobenzidine (DAB) reagent, counterstained with hematoxylin, dehydrated, and mounted.

**RNAseq analysis of lung tissue from the SARS-COV-2 infected mice**

Lung tissue samples were collected and processed in P3 laboratory for safety issues according to the regulation. Total RNA was then extracted and used for the construction of cDNA libraries by an Illumina TruseqTM RNA sample prep Kit. Sequence-by-synthesis single reads of 54-base-length using the Hiseq2000 Truseq SBS Kit (v3-HS, Illumina) were generated on the HiSeq X system. Raw data were collected, and Salmon was used for the quantification of the count and TPM matrix. Subsequently, the tidyverse, clusterProfiler, and GSVA packages of R were used for the downstream analysis. Briefly, the GSVA scores were calculated based on geneset information provided by the msigdbr packages, and the GSVA score for SARS or COVID containing genesets were selected, and the core enriched genes were then extracted as well. All heatmaps or dot plots were created by the pheatmap or ggplot2 packages.

**Statistical analysis**

The data in the figures represent mean ± SD. All data were analyzed using GraphPad Prism 9.0 software. Statistical comparison between different groups was performed using corresponding statistical analysis labeled in figure legends combining several experiments. *P*-values were calculated, and statistical significance was reported as highly significant with ***$P < 0.001$.

**Data availability**

Crystal structures generated during the current study are available in the Protein Data Bank (PDB) under accession codes 8YX2 ($PL^{pro}$ bound to GZNL-P4), 8YX3 ($PL^{pro}$ bound to GZNL-

P28), 8YX4 (PL$^{pro}$ bound to GZNL-P31) and 8YX3 (PL$^{pro}$ bound to GZNL-P35). All other datasets generated and/or analyzed during the current study are available from the corresponding author upon reasonable request.
